# Long-term mitotic DNA damage promotes chromokinesin-mediated missegregation of polar chromosomes in cancer cells

**DOI:** 10.1101/2022.07.22.501103

**Authors:** Marco Novais-Cruz, António Pombinho, Mafalda Sousa, André F. Maia, Helder Maiato, Cristina Ferrás

## Abstract

DNA damage response (DDR) during interphase involves active signalling and repair to ensure genomic stability. However, how mitotic cells respond to DNA damage remains poorly understood. Supported by correlative live-/fixed-cell microscopy analysis we found that mitotic cells exposed to several cancer chemotherapy compounds acquire and signal DNA damage, regardless of how they interact with DNA. In-depth analysis upon long-term DNA damage during mitosis revealed a spindle assembly checkpoint (SAC)-dependent, but DDR-independent, mitotic delay. This delay was due to the presence of misaligned chromosomes that ultimately satisfy the SAC and missegregate, leading to micronuclei formation. Mechanistically, we show that long-term mitotic DNA damage specifically stabilizes kinetochore-microtubule attachments in cancer cells, causing the missegregation of polar chromosomes due to the action of arm-ejection forces by chromokinesins. Overall, these findings unveil that long-term therapeutic DNA damage regimens contribute to genomic instability through a surprising link between the stabilization of kinetochore-microtubule attachments and chromokinesin-mediated missegregation of polar chromosomes in cancer cells.

## Introduction

Genomic DNA is continuously damaged by endogenous and exogenous sources that can generate several types of DNA lesions. To preserve genomic stability in response to DNA damage, cells have evolved a sophisticated system known as the DNA damage response (DDR), which controls DNA repair pathways in coordination with several cell cycle checkpoints (Hustedt and Durocher, 2017; Jackson and Bartek, 2009). Among DNA lesions, double strand breaks (DSBs) are the most deleterious and represent the greatest threat to genomic integrity (Jackson, 2002). Mitotic cells are more vulnerable to DNA damage because most canonical DNA damage repair pathway are inactivated during mitosis. The presence of DSBs in mitosis leads to partial activation of DDR through the recruitment of the MRE11-RAD50-NBS1 (MRN) complex and activation of ataxia telangiectasia mutated (ATM). ATM-mediated phosphorylation of the histone variant H2AX on serine 139 (γH2AX) promotes the recruitment of the mediator of DNA damage checkpoint 1 (MDC1) to damaged sites (Benada et al., 2015; Giunta et al., 2010; van Vugt et al., 2010). However, the mitotic kinases Cdk1 and Plk1 phosphorylate ring finger protein 8 (RNF8) and p53-binding protein 1 (53BP1) to inhibit their recruitment to DSBs and prevent DNA repair, which could otherwise trigger unintended genomic rearrangements, such as telomere fusions (Benada et al., 2015; Lee et al., 2014; Nelson et al., 2009; Orthwein et al., 2014).

Acute exposure to ionizing radiation (IR) during antephase (late G2 to mid prophase) can reverse cell cycle progression, causing cells to return to G2 (Pines and Rieder, 2001). However, acute induction of DNA damage once cells commit to mitosis after nuclear envelope breakdown (NEBD) does not allow cell cycle reversal, but delays mitotic progression in a spindle assembly checkpoint (SAC)-dependent manner (Carlson, 1950; Gomez Godinez et al., 2020; Hayashi et al., 2012; Mikhailov et al., 2002; Silva et al., 2014; Zirkle, 1970). In contrast, acute induction of DSBs during mitosis was found to cause an increase in whole-chromosome segregation errors due to formation of anaphase lagging chromosomes, with no apparent delay in mitotic progression (Bakhoum et al., 2014). These differences might depend on the extent and localization of the DNA damage, as well as on the hyper-stabilization of kinetochore-microtubule attachments due to Aurora A and Plk1 activation upon DNA damage (Bakhoum et al., 2014). Anaphase lagging chromosomes may result in the formation of micronuclei (Soto et al., 2019), which have been recently implicated in genomic instability as critical intermediates of massive chromosome rearrangements associated with chromothripsis (Crasta et al., 2012; Janssen et al., 2011; Shoshani et al., 2021; Zhang et al., 2015).

The combination of DNA-damaging compounds with microtubule-targeting drugs that compromise mitosis are widely used in current cancer chemotherapy regimens (Poruchynsky et al., 2015). Thus clarifying the role of mitotic DNA damage has important clinical implications. Here, we investigated the mechanism by which a broad range of DNA-damaging compounds interfere with mitotic progression and fidelity using a combination of live- and fixed-cell assays, together with specific molecular perturbations. Surprisingly, we found that long-term DNA damage specifically during mitosis interferes with chromosome congression to the spindle equator and induces a SAC-dependent, but DDR-independent, mitotic delay. Importantly, misaligned chromosomes ultimately satisfy the SAC and lead to micronuclei formation. Mechanistically, we show that the combination of the stabilization of kinetochore-microtubule attachments upon mitotic DNA damage and the activity of chromokinesins on chromosome arms leads to the missegregation of polar chromosomes specifically in cancer cells. Taken together, our findings unveil a previously overlooked route by which long-term DNA damage during mitosis contributes to whole-chromosome missegregation. Thus, by driving chromosomal instability, long-term mitotic DNA damage might represent both a threat and an opportunity to chemotherapeutic approaches involving the use of DNA-damaging drugs.

## Results

### The extent of mitotic DNA damage is dependent on the type of DNA lesion

To investigate the impact of DNA damage during mitosis we started by screening a chemical library of 448 cell cycle inhibitors/DNA-damaging compounds using high-content fluorescence microscopy. To determine the respective extent of DNA damage we quantified the phosphorylation levels of histone H2AX on serine 139 [γH2AX; (Rogakou et al., 1998)]. To assist in the identification of mitotic cells, we simultaneously measured the phosphorylation of Histone H3 at serine 10 [pH3S10; (Crosio et al., 2002)]. We used the Z’-factor, a dimensionless, simple statistical coefficient that reflects both the signal dynamic range and variability (Zhang et al., 1999), as a measure of the robustness of our high-throughput assay in identifying mitotic cells with DNA damage (a Z’-factor between 0.5-1 is an excellent indicator of robustness). Accordingly, the Z’-factor between DMSO (negative control) and etoposide (positive control) was 0.76 for mitotic cells (Figure 1A) and 0.60 for interphase cells (Figure S1A). From the screened library, only the compounds that presented a percentage of cells with γH2AX higher than the median plus three standard deviations (3SD) of the negative control were considered as hits. Based on this criterion, we identified 73 hits that induced DNA damage in mitotic cells, 10 of which did not induce detectable DNA damage (or the damage was rapidly repaired) in interphase cells (Figure 1A-C). In parallel, we identified 95 compounds that induced DNA damage in interphase cells (Figure S1A), 32 of which did not cause detectable DNA damage in mitotic cells (Figure 1C).

**Figure 1.**
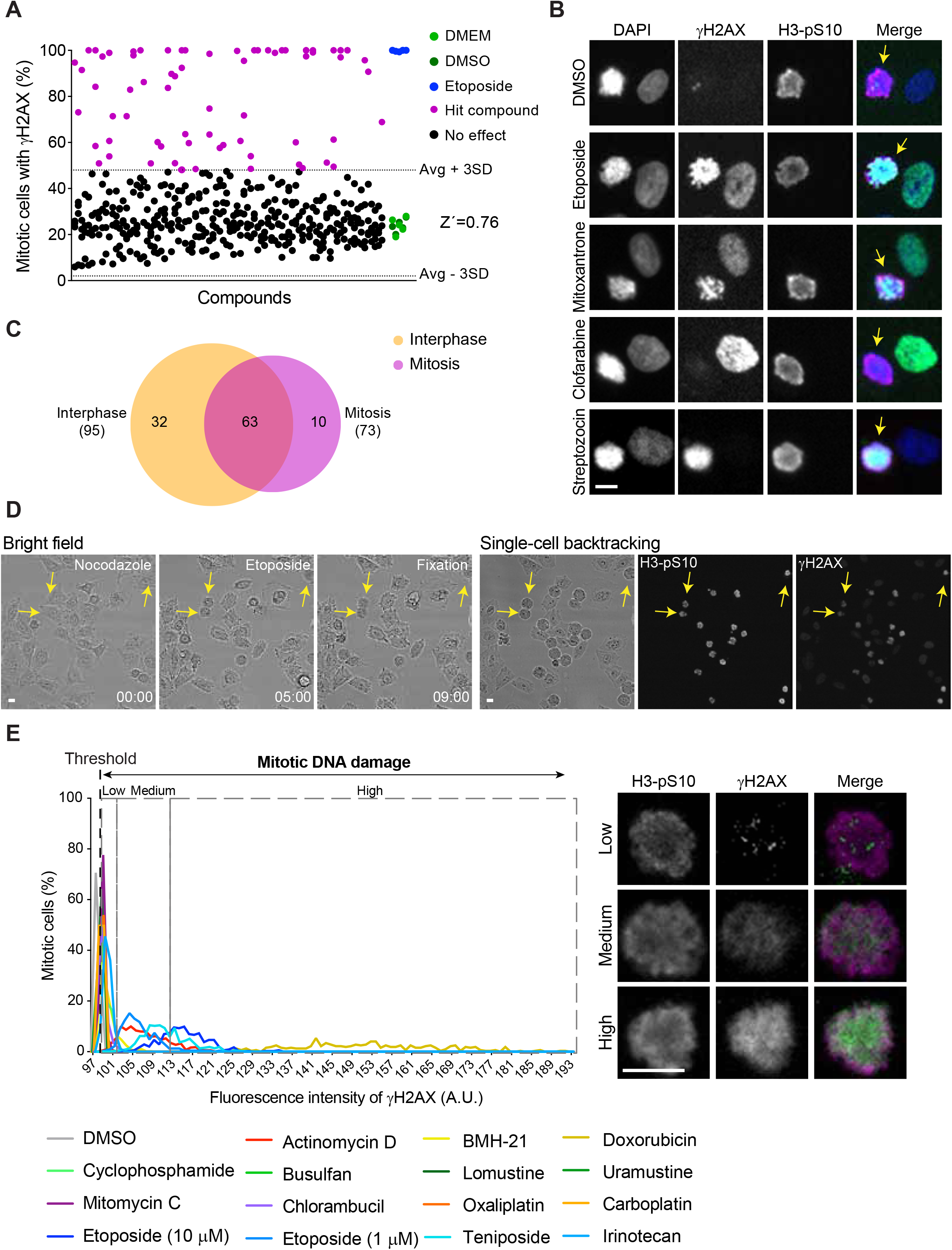
HeLa cells acquire specifically different levels of DNA damage in mitosis. **(A)** Percentage of mitotic HeLa cells with γH2AX positive after 4 h treatment with chemical library of 448 cell cycle inhibitors/DNA-damaging compounds. DMEM and DMSO (green dots) and etoposide (blue dots) were used as negative and positive controls. Each dot represents a compound. The thresholds (dash lines) were defined as mean (Avg) plus or minus three times standard deviation (3SD) from negative controls. Compounds above the threshold were considered hits (magneta dots). **(B)** Representative images of negative (DMSO) and positive (etoposide) control and hits compounds (mitoxantrone, clofarabine and streptozocin). Scale bar = 10 μm. **(C)** Venn diagram of identified hits in interphase and mitotic HeLa cells from 448 tested compounds. **(D)** *(left side)* Selected time frames from bright field microscopy of HeLa cells treated with nocodazole and etoposide. Images were acquired every 20 min. Time = h:min. *(right side)* Immunofluorescence imagens of the last time frame (09:00) from bright field microscopy with the indicated antibodies. Arrows highlight examples of cells already committed to mitosis when exposed to the drug. Scale bar = 10 μm. **(E)** *(left side)* Histograms of selected mitotic DNA-damaging compounds (n = 14), showing fluorescence intensity of γH2AX in mitotic HeLa cells. The threshold (black dash line) was defined as the lowest fluorescence intensity detected for γH2AX foci. The different categories (low, medium, high; dashed gray squares) were defined based on intensity levels of γH2AX. *(right side)* Representative imagens of different intensity levels of γH2AX in mitotic HeLa cells. Scale bar = 10 μm.

Next, to investigate whether a subset of the identified compounds is able to induce DNA damage when cells were already committed to mitosis, we implemented a correlative live-cell microscopy and quantitative immunofluorescence analysis assay using γH2AX. Briefly, human HeLa cells were tracked by time-lapse bright field microscopy after treatment with the microtubule depolymerizing drug nocodazole. After 5 h in nocodazole, we treated cells for 4 h with 14 of the 73 mitotic DNA-damaging compounds that are currently in clinical use and, proceeded to fixation. Subsequently, the levels of γH2AX were analysed by fluorescence microscopy and the history of each cell was inferred by back-tracking using the corresponding live-cell data, confirming that DNA damage was inflicted when cells were already committed to mitosis (Figure 1D,E). We found that lesions caused either by DNA-alkylating compounds (cyclophosphamide, busulfan, lomustine, uramustine), DNA-crosslinking agents (mitomycin c, chlorambucil) or DNA adduct-inducing drugs (oxaliplatin, carboplatin) result in low levels of mitotic DNA damage. Except for BMH-21 and irinotecan that induced low levels of mitotic DNA damage, treatment with DNA-intercalating compounds (actinomycin D and doxorubicin), or drugs that generate single- or double-strand breaks (etoposide and teniposide), resulted in higher levels of mitotic DNA damage (Figure 1E). Western blot analysis of mitotic cells treated with the selected DNA-damaging compounds confirmed these results (Figure S1B). Taken together, we conclude that HeLa cells may acquire DNA damage specifically during mitosis and the extent of the damage is dependent on the type of DNA lesion.

### Long-term mitotic DNA damage delays anaphase onset due to the formation of polar chromosomes that result in micronuclei

To investigate the impact of long-term mitotic DNA damage for the progression and exit of mitosis, we used live-cell fluorescence microscopy to directly determine the mitotic duration and respective cell fate. To assure the induction of long-term DNA damage during mitosis we implemented a nocodazole treatment/washout protocol. Briefly, human HeLa cells were partially synchronized in mitosis upon treatment with the microtubule depolymerizing drug nocodazole for 3 h. Subsequently, selected representative compounds that induce distinct DNA lesions (lomustine, mitomycin C, carboplatin, and etoposide) were added for 4 h. After this period, nocodazole was washed out and the cells allowed to progress through mitosis in the presence of the respective DNA-damaging compounds. Under these conditions, control DMSO-treated cells took 128 ± 85 min (mean ± SD) to enter anaphase after nocodazole washout and 66% of the cells divided without any detectable segregation defect (Figure 2A-C and Figure S2). In contrast, among other defects, treatment with any of the DNA-damaging compounds caused a significant mitotic delay due to the presence of misaligned chromosomes near the spindle poles (from here on, referred as polar chromosomes) (Figure 2A-C and Figure S2). Despite the observed mitotic delay, most cells were able to enter anaphase, some of which (11-18%, depending on the condition) without ever completing congression of polar chromosome(s) that ultimately gave rise to micronuclei (Figure 2C).

**Figure 2.**
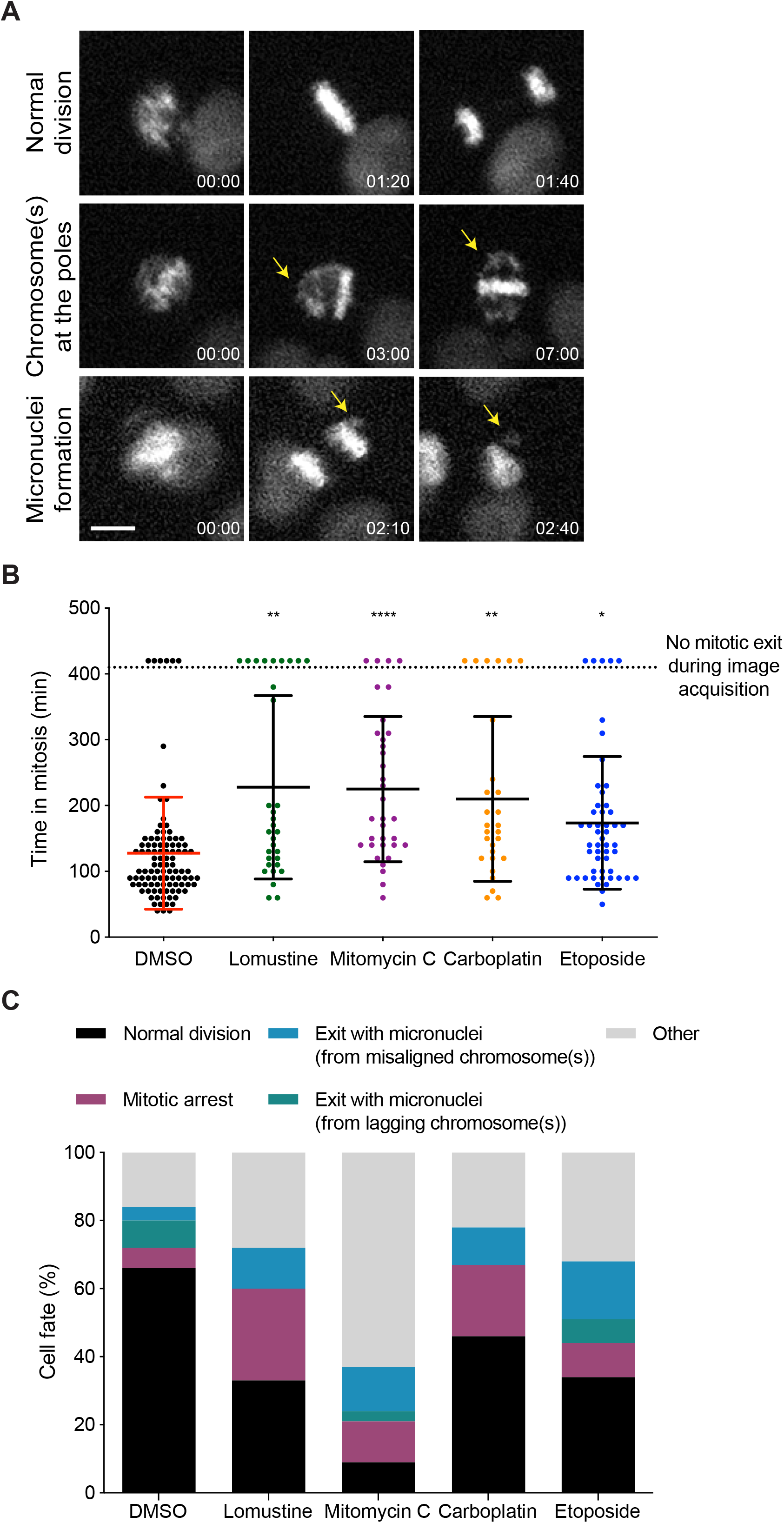
Long-term mitotic DNA damage induces a mitotic delay due to the presence of polar chromosomes, leading to micronuclei formation. **(A)** Representative images of time-lapse sequences illustrating the three main mitotic phenotypes observed. Arrows highlight chromosomes at the poles or micronuclei formation. Pixels were saturated for optimal visualization of misaligned chromosomes and micronuclei. Images were acquired every 10 min. Scale bar = 10 μm. Time = h:min. **(B)** Mitotic duration between nocodazole washout and anaphase onset in mitotic HeLa cells treated with DMSO or selected mitotic DNA-damaging compounds. Each data point corresponds to one cell. (DMSO, 127.6 ± 85.1 min, n = 101; Lomustine, 227.8 ± 139.3 min, n = 32; Mitomycin C, 225.0 ± 110.3 min, n = 34; Carboplatin, 210.0 ± 125.2 min, n = 28; Etoposide, 173.7 ± 100.8 min, n = 51; mean ± SD; * p<0.05, ** p<0.01, ****p≤0.0001 relative to control, multiple comparison Kruskal-Wallis test). **(C)** Cell fate of mitotic HeLa cells treated with the same drugs as in B.

To test whether the induction of long-term DNA damage in prometaphase cells persists throughout mitosis, we fixed cells 70 min after nocodazole washout. We found that γH2AX foci remained on chromosomes in cells that have reached metaphase and entered anaphase (Figure S3A,B). These results indicate that damaged sites are recognized but not repaired during mitosis, in agreement with previous observations (Ciccia and Elledge, 2010; Giunta et al., 2010; Giunta and Jackson, 2011). Taken together, these experiments show that, regardless of the type of DNA lesion, long-term mitotic DNA damage persists throughout mitosis and promotes micronuclei formation from polar chromosomes.

### The mitotic delay observed upon long-term DNA damage is exclusively mediated by the SAC

It was previously proposed that DDR and the SAC cooperate to delay mitotic progression in the presence of DNA damage in budding yeast (Kim and Burke, 2008). However, whether this is the case in human cells remains unclear. To clarify this, we performed live-cell imaging to monitor mitotic duration and cell fate upon long-term mitotic DNA damage and subsequent SAC abrogation with the Mps1 inhibitor MPS1-IN-1. We found that treatment with either MPS1-IN-1 alone or in combination with mitotic DNA-damaging compounds caused a fast mitotic exit in nocodazole-arrested HeLa cells (Figure 3A,B). This phenotype was fully reverted by treating cells with the proteasome inhibitor MG132 (Figure 3A,B). These results indicate that the observed mitotic delay upon long-term DNA damage is exclusively mediated by the SAC.

**Figure 3.**
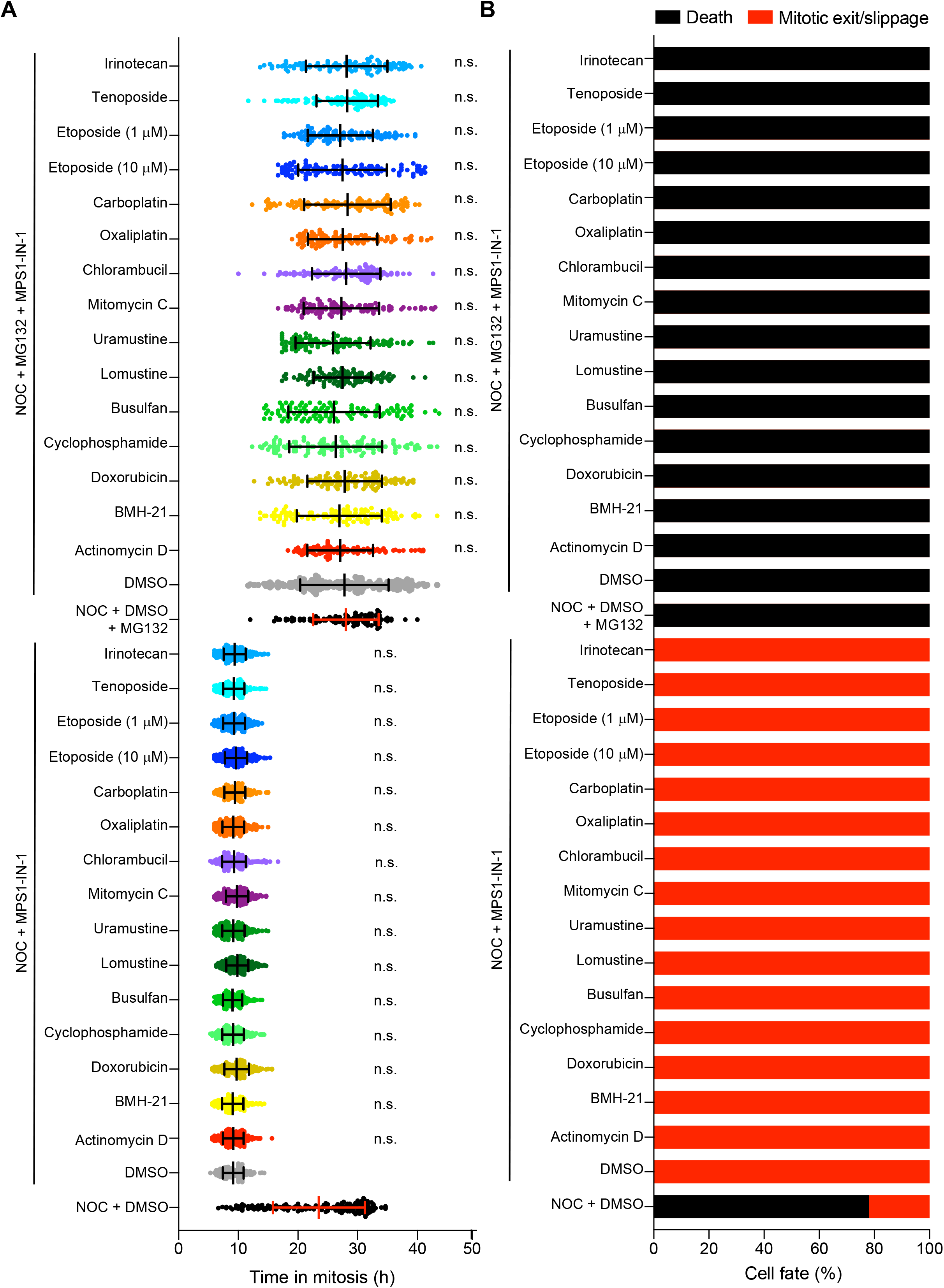
DNA damage response does not contribute to arrest cells in mitosis. **(A)** Mitotic duration of the mitotic arrest of HeLa cells treated with nocodazole and Mps1 inhibitor (MPS1-IN-1) in combination with selected mitotic DNA-damaging compounds with or without MG132. Each data point corresponds to one cell. The results were expressed as mean ± SD; n.s.: not significant relative to control, multiple comparison Kruskal-Wallis test. **(B)** Cell fate of mitotic HeLa cells treated with the same drugs as in A.

### Long-term mitotic DNA damage leads to the formation of polar chromosomes that eventually form stable kinetochore-microtubule attachments

To determine the status of kinetochore-microtubule attachments on polar chromosomes that form upon long-term mitotic DNA damage, we investigated the localization of the SAC protein Mad1, which accumulates at unattached kinetochores delaying anaphase onset (Musacchio, 2015). In this case, cells were fixed 70 min after nocodazole washout, a time window in which we detected polar chromosomes with or without DNA damage. In striking contrast to control DMSO-treated cells where Mad1 signal was mostly detected on both kinetochores, mitotic DNA damage significantly increased the frequency of polar chromosomes in which only the most distal kinetochore remained Mad1-positive, indicating the establishment of stable monotelic attachments (Figure 4A,B). Similar findings were observed after mitotic arrest/washout with the kinesin-5 inhibitor S-trityl-L-cysteine (STLC) upon long-term DNA damage, ruling out a nocodazole/microtubule specific effect (Figure S4A,B). Importantly, the observed Mad1 asymmetry after mitotic DNA damage was not due to differences in the total levels of Mad1 protein (Figure 4C) and phenocopied the behaviour observed upon microtubule stabilization with 20 nM taxol (Figure 4D,E). These results suggest that long-term mitotic DNA damage promotes the formation of polar chromosomes that eventually form stable kinetochore-microtubule attachments.

**Figure 4.**
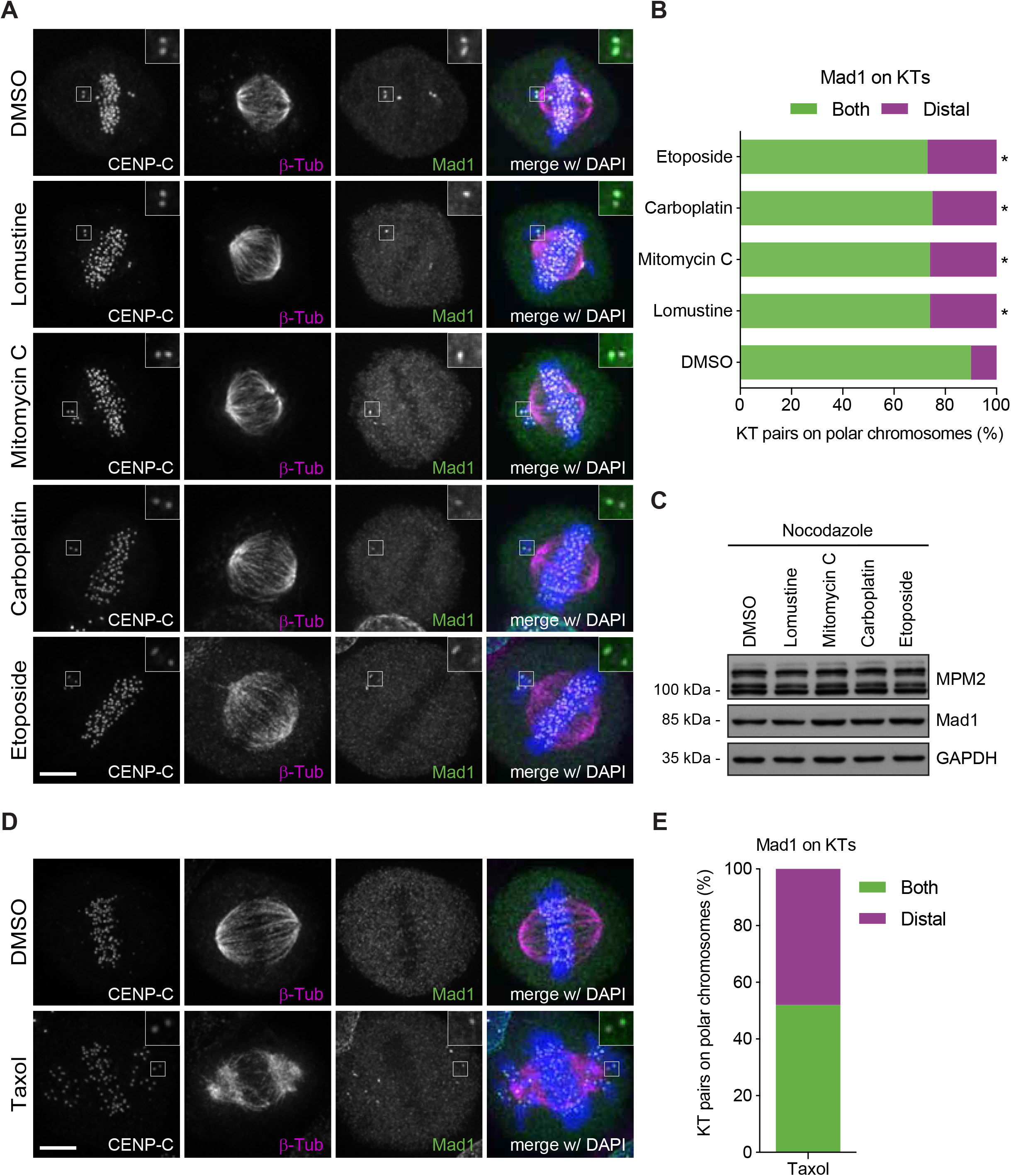
Long-term mitotic DNA promotes the formation of polar chromosomes that form stable monotelic attachments. **(A)** Representative immunofluorescence images of HeLa cells treated with nocodazole and either DMSO or different DNA-damaging compounds during 4 h, followed by nocodazole washout and fixation after 70 minutes. Cells were immunostained with anti-CENP-C (gray), anti-β-tubulin antibody (magneta), anti-Mad1 antibody (green) and DAPI (blue). Scale bar = 5 μm. **(B)** Quantification of Mad1 detection on kinetochores pairs from polar chromosomes under different conditions presented in A. **(C)** Western blot analysis of Mad1 levels from mitotic HeLa cell extracts. Mitotic extracts were derived from isolated mitotic cells treated with DMSO, Etoposide, Lomustine, Mitomycin C or Carboplatin. GAPDH was used as loading control. Approximate molecular weights are shown on the left. **(D)** Representative immunofluorescence images of HeLa cells treated with DMSO or taxol during 2 h. Cells were immunostained with anti-CENP-C (gray), anti-β-tubulin antibody (magneta), anti-Mad1 antibody (green) and DAPI (blue). Scale bar = 5 μm. **(E)** Quantification of Mad1 detection on kinetochores pairs from polar chromosomes under different conditions presented in D.

### Long-term mitotic DNA damage increases kinetochore-microtubule attachment stability in cancer cells

To directly investigate the effect of long-term mitotic DNA damage on kinetochore-microtubule attachment stability, we determined kinetochore-microtubule half-life upon a nocodazole shock in HeLa cells (Warren et al., 2020). In agreement with previous measurements using fluorescence dissipation after photoactivation of PA-GFP-α -tubulin (Bakhoum et al., 2014), this assay revealed that kinetochore-microtubules were more resistant to nocodazole-induced depolymerization upon long-term mitotic DNA damage, as determined by a significant increase in microtubule half-life from 2.34 min without mitotic DNA damage, to 3.90 min upon mitotic DNA damage (Figure 5A,B).

**Figure 5.**
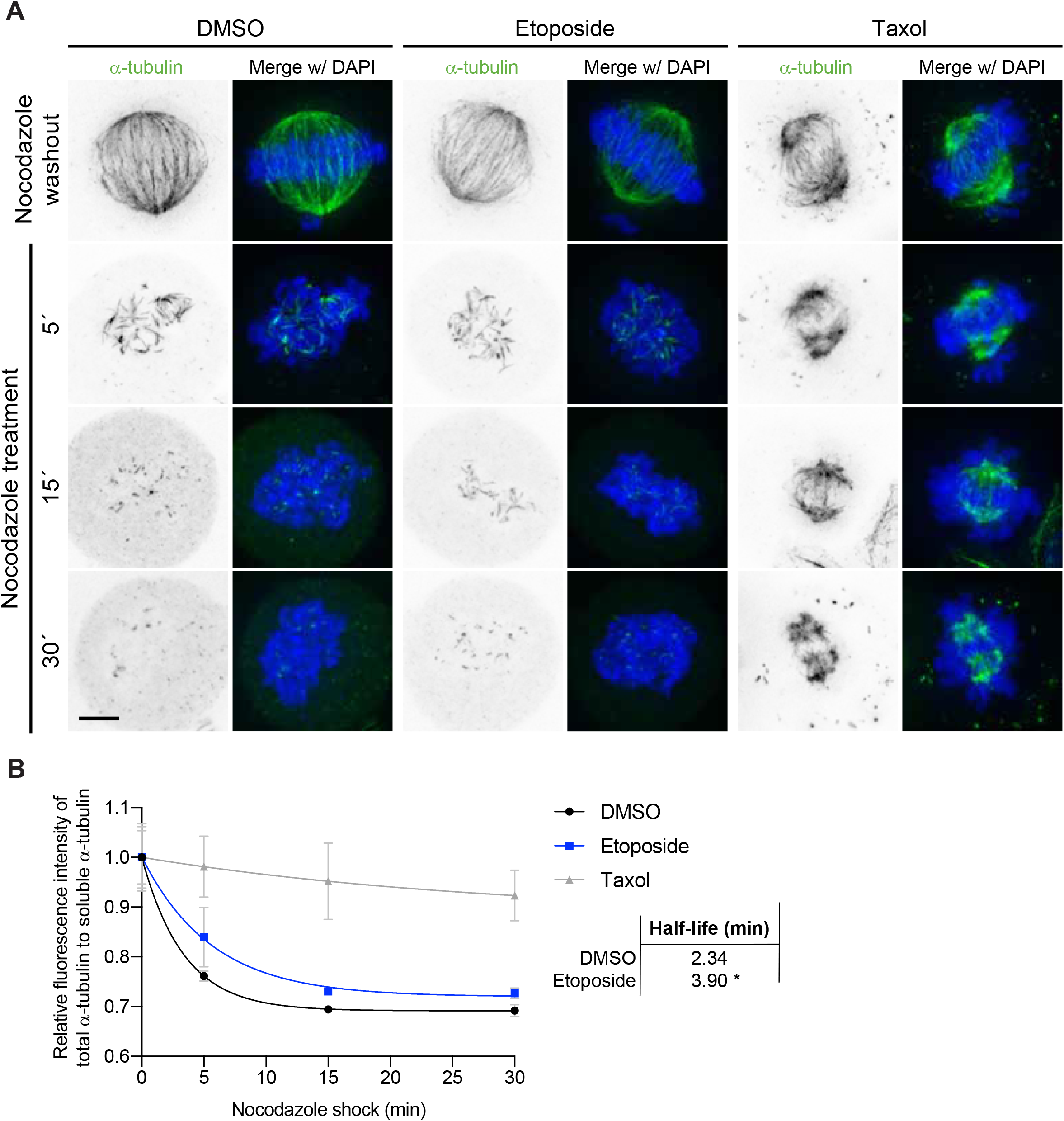
Long-term mitotic DNA damage increases kinetochore-microtubule attachment stability in HeLa cells. **(A)** Representative immunofluorescence images of HeLa cells treated with nocodazole and either DMSO or etoposide during 4 h, followed by nocodazole washout. After 70 minutes, a nocodazole shock was applied and the cells were fixed after 5, 15 and 30 minutes. Cells were immunostained with anti-β-tubulin antibody (inverted grey scale) and DAPI (blue). Scale bar = 5 μm. Taxol was used as a positive control. **(B)** Normalized ratio of total α-tubulm soluble α-tubulin fluorescence intensity over time. The kinetics of fluorescence decay after nocodazole shock were determined as a readout of resistance to microtubule depolymerization. Whole lines show single exponential fitting curve. Each data point represents the mean ± SD; * p<0.05 relative to control, extra sum-of-square F test.

To test whether the observed stabilization of kinetochore-microtubule attachments upon long-term mitotic DNA damage is a common feature between cancer and non-cancer cells, we extended our analysis to measure microtubule half-life in both transformed chromosomally unstable U2OS cells and hTERT-immortalized, but not transformed, near-diploid RPE-1 cells. Similar to HeLa cells, we found that etoposide induced mitotic DNA damage in U2OS cells (Figure S5) and significantly increased the frequency of polar chromosomes with single unattached kinetochores (as inferred by Mad1 immunofluorescence detection) (Figure S6A,B). Most relevant, U2OS cells showed a significant increase in microtubule half-life upon long-term mitotic DNA damage (Figure S6C,D). In sharp contrast, for an equivalent time window, all polar chromosomes complete congression after long-term mitotic DNA damage and nocodazole washout in RPE-1 cells (our unpublished observations), which showed no detectable differences in kinetochore-microtubule half-life relative to DMSO-treated controls (Figure S7A-C). Overall, these results indicate that the formation of polar chromosomes due to the stabilization of kinetochore-microtubule attachments upon long-term mitotic DNA damage is likely exclusive to cancer cells.

### Long-term mitotic DNA damage causes missegregation of polar chromosomes due to the action of arm-ejection forces by chromokinesins

The kinetochore-microtubule attachment status of polar chromosomes depends on the coordinated activities of kinetochore- and arm-associated motor proteins that ultimately determine the proximity to the spindle pole where the microtubule-destabilizing activity of Aurora A is higher (Barisic et al., 2014; Barisic and Maiato, 2015; Ye et al., 2015). For instance, inhibition of the microtubule plus-end-directed motor CENP-E at kinetochores generates polar chromosomes in the immediate vicinity of the spindle poles with both kinetochore pairs unattached, due to the microtubule minus-end-directed motor activity of Dynein at kinetochores (Barisic et al., 2014; Maia et al., 2010; McEwen et al., 2001; Putkey et al., 2002). In contrast, simultaneous inhibition of both CENP-E and Dynein motor activities at kinetochores, results in the cortical ejection of polar chromosomes away from high Aurora A activity at the poles due to the action of the chromokinesins Kid/kinesin-10 and Kif4a/kinesin-4 on chromosome arms (Almeida and Maiato, 2018) resulting in the stabilization of monotelic kinetochore-microtubule attachments (Barisic et al., 2014; Barisic and Maiato, 2015; Cane et al., 2013; Drpic et al., 2015; Ye et al., 2015). Given that long-term mitotic DNA damage results in the formation of polar chromosomes that establish stable monotelic attachments, we investigated whether this stabilizing effect was dependent on chromokinesin-mediated ejection of polar chromosomes away from high Aurora A activity at the poles. Interestingly, it has been shown that, upon ionizing radiation, Kif4a is recruited to damaged DNA sites in interphase cells (Wu et al., 2008). However, the total levels of Kif4a (and Kid) on chromosome arms were not altered upon long-term mitotic DNA damage (Figure 6A-D). Moreover, neither Aurora A activity at spindle poles, nor Aurora B centromeric activity on polar chromosomes, was affected upon long-term mitotic DNA damage (Figure S8A-D). Nevertheless, co-depletion of both Kid and Kif4a upon mitotic long-term DNA damage significantly reverted the formation of polar chromosomes with stable monotelic attachments (Figure 6E-G). Altogether, these results support a model in which long-term mitotic DNA damage specifically stabilizes kinetochore-microtubule attachments in cancer cells, causing the missegregation of polar chromosomes due to the action of arm-ejection forces by chromokinesins.

**Figure 6.**
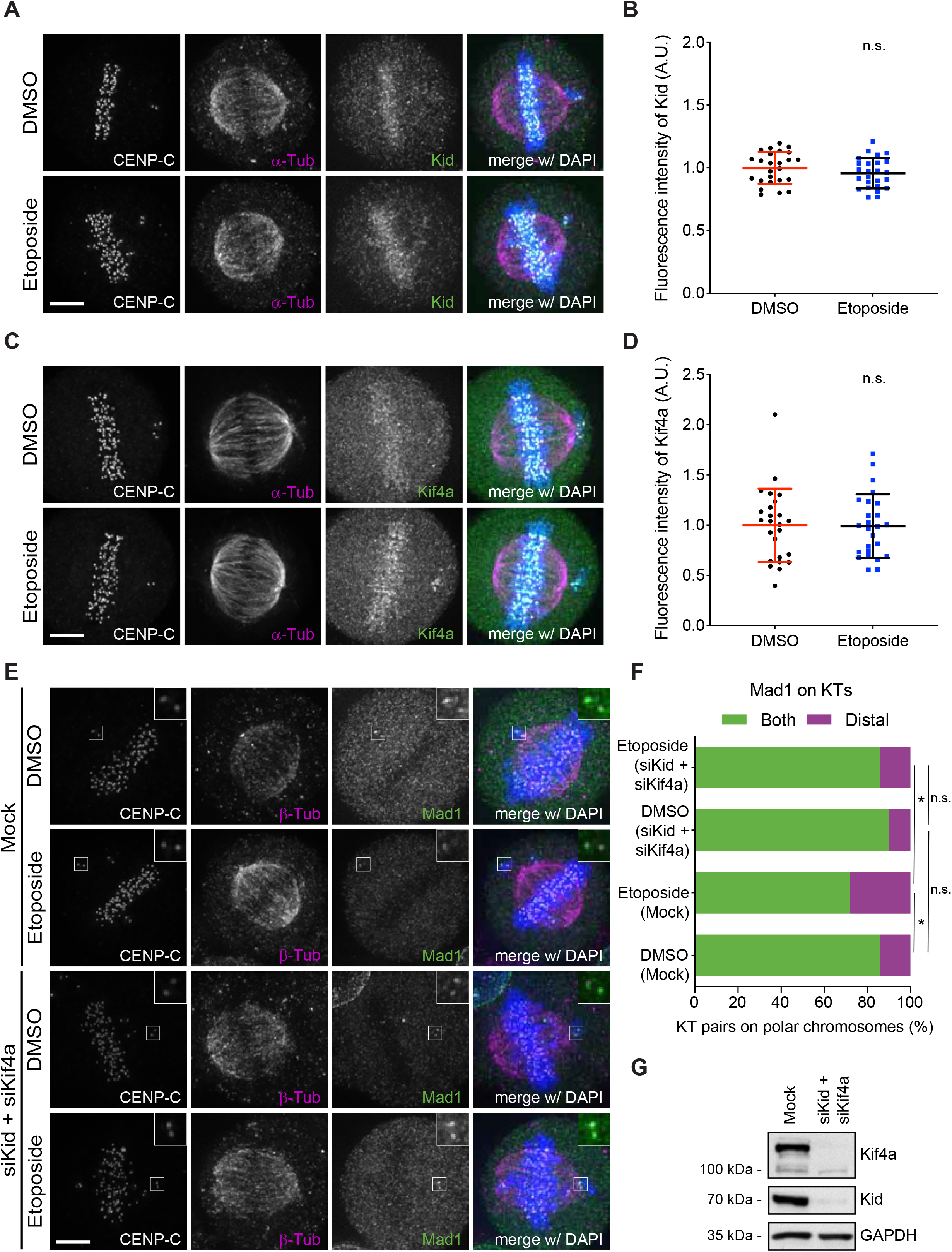
Long-term mitotic DNA damage promotes missegregation of polar chromosomes due to the chromokinesins activity. **(A)** Representative immunofluorescence images of HeLa cells treated with nocodazole and either DMSO or etoposide during 4 h, followed by nocodazole washout and fixation after 70 minutes. Cells were immunostained with anti-CENP-C (gray), anti-α-tubulin antibody (magneta), anti-Kid antibody (green) and DAPI (blue). Scale bar = 5 μm. **(B)** Normalized fluorescence intensity of Kid. Each dot represents an individual cell. (DMSO, 1.00 ± 0.13, n = 25; Etoposide, 0.96 ± 0.12, n = 25; mean ± SD; n.s.: not significant relative to control, t test). **(C)** Representative immunofluorescence images of HeLa cells treated with nocodazole and either DMSO or etoposide during 4 h, followed by nocodazole washout and fixation after 70 minutes. Cells were immunostained with anti-CENP-C (gray), anti-γH2AX antibody (magneta), anti-Kif4a antibody (green) and DAPI (blue). Scale bar = 5 μm. **(D)** Normalized fluorescence intensity of Kif4a. Each dot represents an individual cell. (DMSO, 1.00 ± 0.36, n = 25; Etoposide, 0.99 ± 0.32, n = 25; mean ± SD; n.s.: not significant relative to control, Mann-Whitney Rank Sum test). **(E)** Representative immunofluorescence images of control and Kid/Kif4a-depleted HeLa cells treated with nocodazole and either DMSO or etoposide during 4 h, followed by nocodazole washout and fixation after 70 minutes. Cells were immunostained with anti-CENP-C (gray), anti-β-tubulin antibody (magneta), anti-Mad1 antibody (green) and DAPI (blue). Scale bar = 5 μm. **(F)** Quantification of Mad1 detection on kinetochores pairs from polar chromosomes under different conditions presented in E. **(G)** Western blot analysis of HeLa cell extracts after RNAi with the indicated antibodies. GAPDH was used as loading control. Approximate molecular weights are shown on the left.

## Discussion

Several DNA-damaging compounds are currently used in combination with microtubule-targeting drugs that compromise mitosis in several cancer treatment regimens. Thus, understanding the consequences of long-term DNA damage during mitosis remains a fundamental question in cell biology, with strong clinical implications. Previous studies suggested that DNA damage in mitosis is signalled but not repaired. Here we show that long-term DNA damage induced after cells commit to mitosis (i.e. prometaphase) remains throughout metaphase and anaphase, supporting the idea of partial activation of DDR in mitosis. Interestingly, even without additional external DNA-damaging sources, a prolonged mitotic arrest induced by microtubule poisons (e.g. nocodazole) was shown to promote the accumulation of DNA damage and to delay mitotic exit after drug washout (Dalton et al., 2007; Smits et al., 2000). However, in contrast with budding yeast cells that undergo a closed mitosis (Kim and Burke, 2008), a partial activation of DDR is insufficient to arrest human cancer cells in mitosis upon long-term DNA damage. Instead, our results indicate that the SAC is the only mechanism that is required to delay mitosis in the presence of long-term mitotic DNA damage. These findings are consistent with a previous work that showed that acute mitotic DNA damage induced by laser microsurgery and topoisomerase 2 inhibitors in human cells during late prophase results in the formation of few unattached kinetochores that delay cells in metaphase in a SAC-dependent manner (Mikhailov et al., 2002). However, much to our surprise, long-term mitotic DNA damage resulted in a SAC-dependent delay of mitotic exit due to the formation of polar chromosomes. In many such cases, cells end up satisfying the SAC, entering anaphase without ever completing congression of polar chromosomes to the spindle equator. We show that long-term mitotic DNA damage specifically stabilizes kinetochore-microtubule attachments in cancer cells, causing the missegregation of polar chromosomes due to the action of arm-ejection forces by chromokinesins, and leading to the formation of micronuclei (Figure 7).

**Figure 7.**
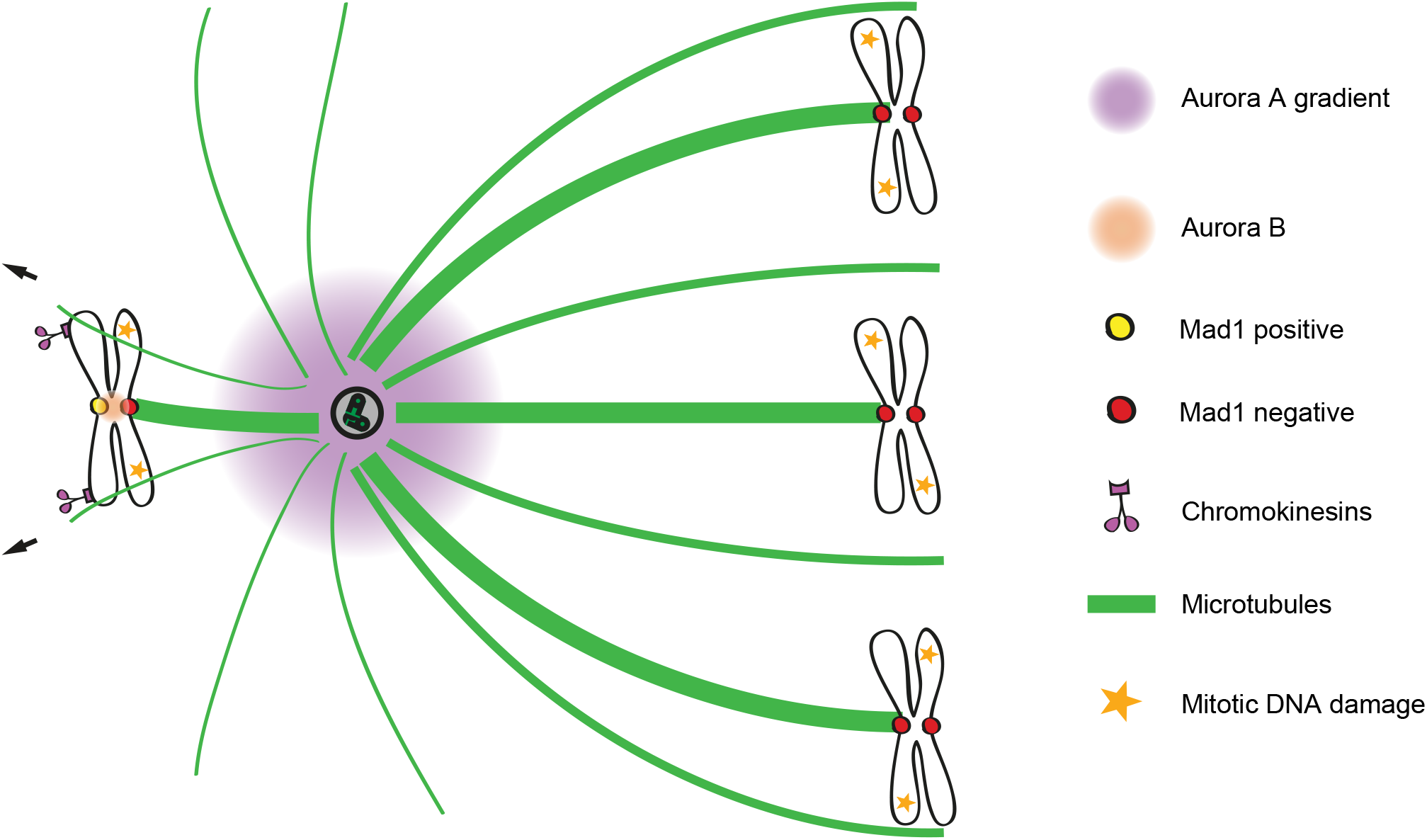
Proposed model for missegregation of polar chromosomes upon long-term mitotic DNA damage. The induction of long-term DNA damage during mitosis stabilizes kinetochore-microtubule attachments, causing the missegregation of polar chromosomes due to the action of arm-ejection forces by chromokinesins.

How DNA damage stabilizes kinetochore-microtubule attachments remains mysterious. Curiously, the chromokinesin Kif4a has been shown to be recruited to DNA-damage sites during interphase (Wu et al., 2008) and enhanced chromokinesin activity has been shown to promote the stabilization of kinetochore-microtubule attachments in *Drosophila* cells (Cane et al., 2013; Drpic et al., 2015). While we cannot exclude that this is also the case for mitotic DNA damage, we were unable to detect any significant enrichment of Kif4a or Kid on DNA-damaged chromosomal sites, leaving open the possibility that mitotic DNA damage might promote chromokinesin activity on chromosome arms. In another study acute mitotic DNA damage was proposed to increase kinetochore-microtubule attachment stability through activation of Aurora A and Plk1 kinases (Bakhoum et al., 2014). However, in this case, increased kinetochore-microtubule attachment stability due to mitotic DNA damage induced the formation of lagging chromosomes during anaphase, which might result in the formation of micronuclei. One possibility supported by our data is that Aurora A activity is not altered in the case of long-term DNA damage in cells delayed in mitosis for several hours. Because most anaphase lagging chromosomes in normal and cancer cells rarely form micronuclei and were recently shown to be actively corrected by an anaphase surveillance mechanism involving Aurora B activity at the spindle midzone (Orr et al., 2021; Sen et al., 2021), the presence of misaligned chromosomes that eventually satisfy the SAC upon long-term DNA damage might represent a higher risk for micronuclei formation and consequent genomic instability due to chromothripsis. Indeed, we recently found that systematic perturbation of kinetochore-microtubule attachments caused human cancer cells to enter anaphase after a delay with chronically misaligned chromosomes that eventually satisfy the SAC and missegregate, leading to the formation of micronuclei (Gomes et al., 2022). Thus, chronically misaligned chromosomes may represent a previously overlooked mechanism driving chromosomal/genomic instability during cancer cell division. This can be seen both as a threat and an opportunity to chemotherapeutic approaches involving the combined use of DNA-damaging compounds with microtubule-targeting drugs that compromise mitosis. On one hand, these combinatorial therapies might result in enhanced cytotoxicity due to a synergistic effect caused by DNA damage and chromosome missegregation. On the other hand, this might contribute to cancer cell adaptation and rapid evolution which might contribute to therapeutic resistance. The results provided in the present study justify further and careful scrutiny of the pros and cons of combinatorial therapies involving DNA-damaging and mitotic drugs in the context of specific cancers.

## Materials and Methods

### Reagents and Tools table

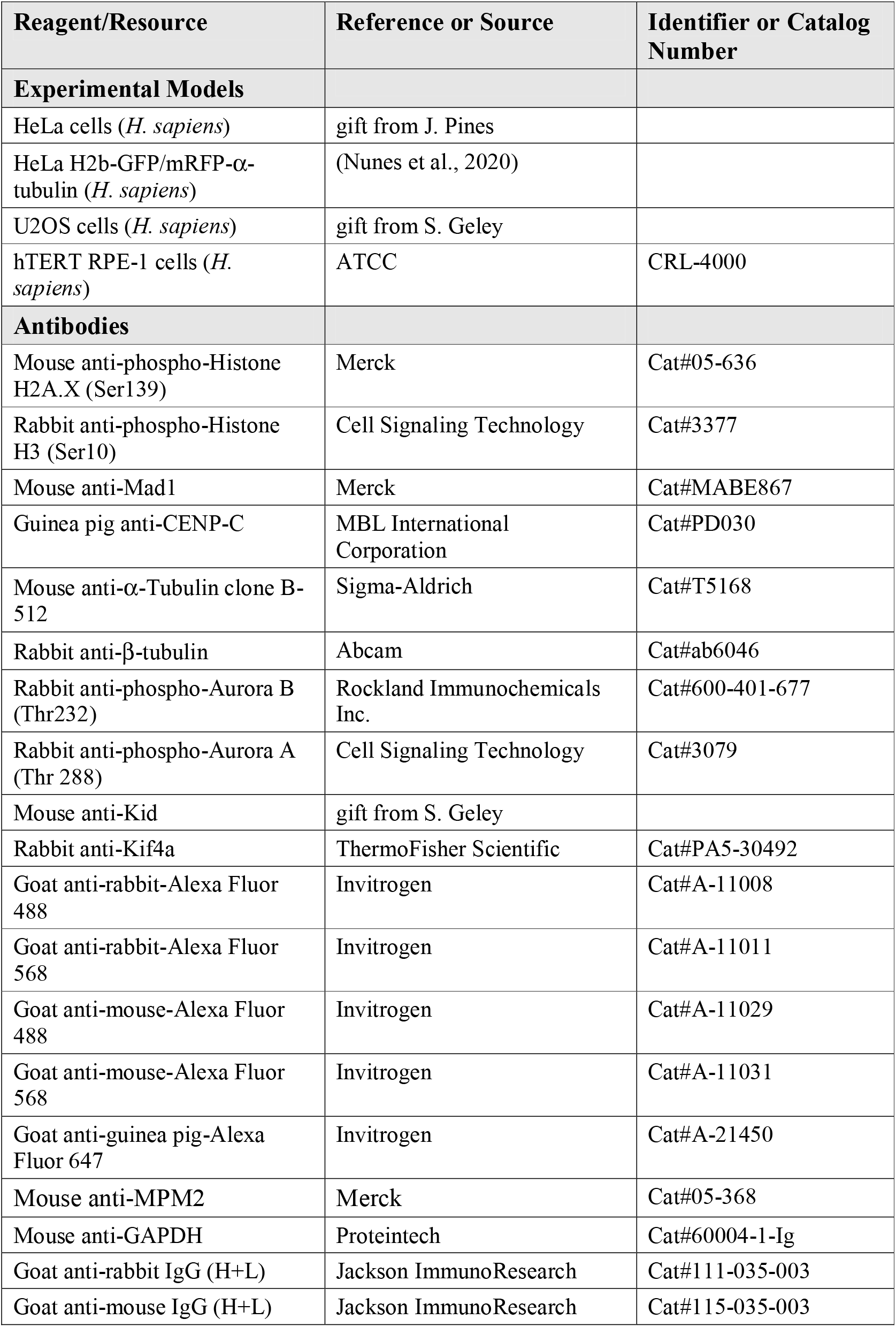
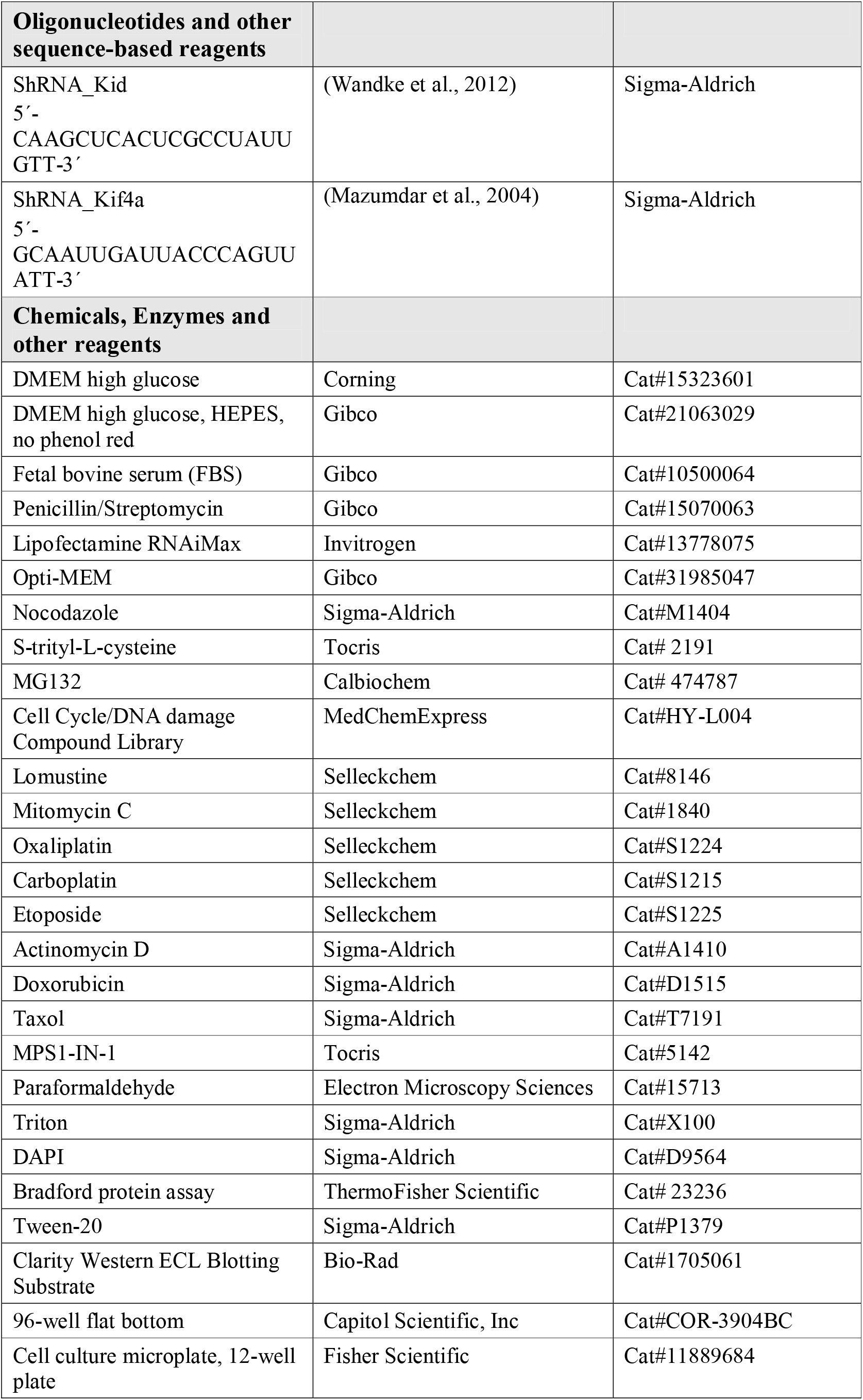
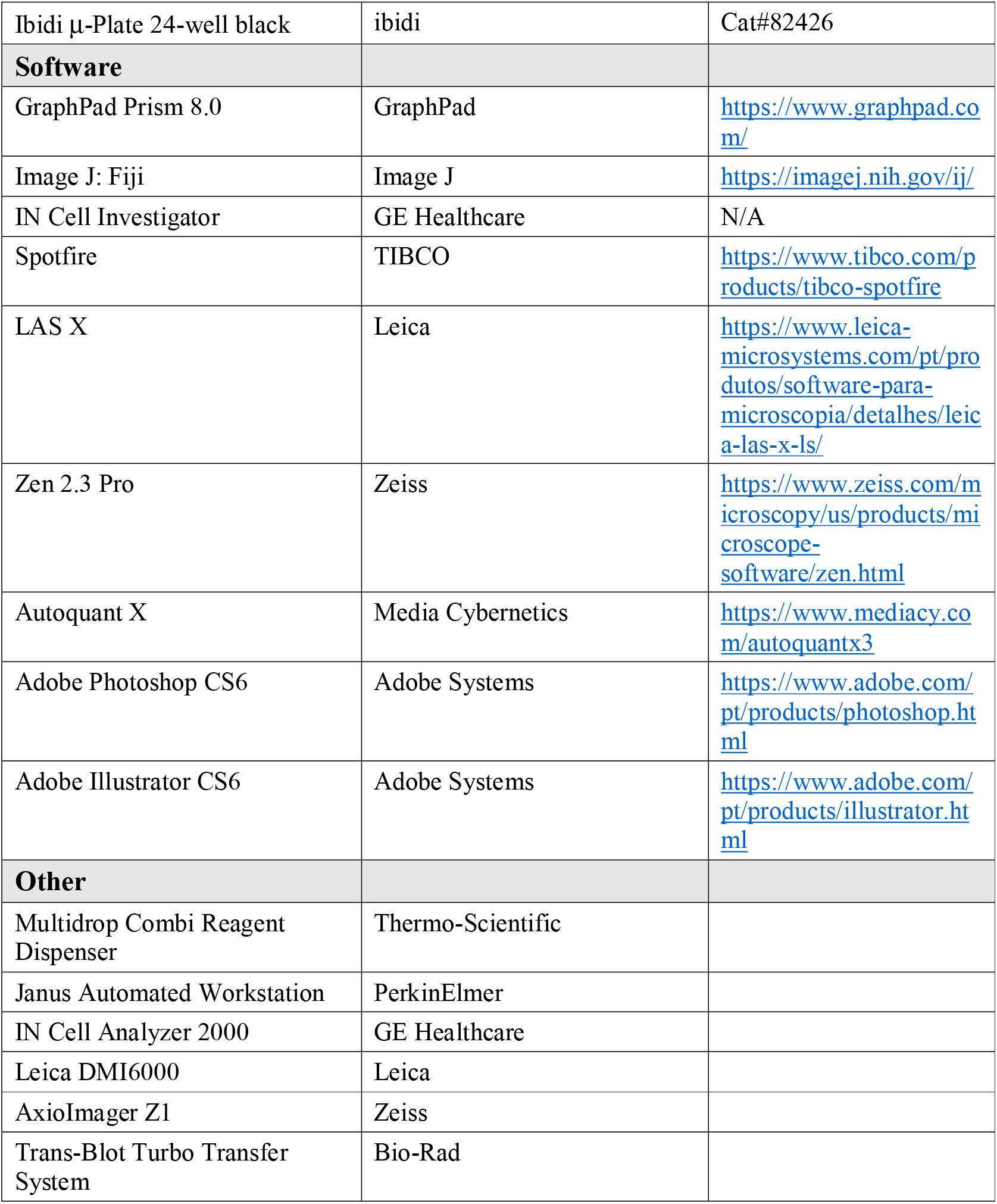

### Cell Culture and siRNA transfections

HeLa parental and U2OS parental were kindly provided by J. Pines (Cancer Research Institute, London, UK) and S. Geley (Innsbruck Medical University, Innsbruck, Austria), respectively. HeLa cells stably expressing GFP-H2B/mRFP-α-tubulin were previously generated in (Nunes et al., 2020). All cell lines, including hTERT RPE-1 parental (ATCC), were grown in Dulbecco’s Modified Eagle Medium (DMEM; Corning) supplemented with 10% fetal bovine serum (FBS; Gibco). All cell lines were grown with penicillin/streptomycin (100 IU/mL and 100 μg/mL; Gibco) at 37°C in a humidified incubator with 5% CO_2_. To performed siRNA experiments, HeLa cells were plated onto 22×22 mm no. 1.5 glass coverslips (Corning) in DMEM supplemented with 5% FBS. siRNA transfection was performed using Lipofectamine RNAiMax (Invitrogen) and 50 nM of the respective siRNA, diluted in serum-free medium (Opti-MEM; Gibco). Mock transfection was used as control. The cells were analysed 72 hr after depletion. Depletion efficiency was evaluated by western blot. The following target sequences were used:

Kid, 5’-CAAGCUCACUCGCCUAUUGTT-3’;

Kif4a, 5’-GCAAUUGAUUACCCAGUUATT-3’;

### Drug Treatments

Microtubule depolymerization was induced by Nocodazole (Sigma-Aldrich) at 3.3 μM for 3-5 h, according to the experiment. HeLa cells were also blocked in mitosis with 10 μM S-trityl-L-cysteine (STLC; Tocris). Mitotic arrest at metaphase or proteasome inhibition was obtained using 5 μM MG132 (Calbiochem). To induce different type of DNA lesions, the following drugs were used for 4 h: 10 μM Cyclophosphamide (MedChemExpress), 10 μM Busulfan (MedChemExpress), 10 μM Lomustine (Selleckchem), 10 μM Uramustine (MedChemExpress), 10 μM Mitomycin C (Selleckchem), 10 μM Chlorambucil (MedChemExpress), 10 μM Oxaliplatin (Selleckchem), 10 μM Carboplatin (Selleckchem), 10 μM or 1 μM Etoposide (Selleckchem), 10 μM Teniposide (MedChemExpress), 10 μM Irinotecan (MedChemExpress), 10 μM BMH-21 (MedChemExpress), 8 μM Actinomycin D (Selleckchem) and 0.5μM Doxorubicin (Sigma-Aldrich). To stabilize microtubules 20 nM Taxol (Sigma-Aldrich) was added to the cell culture media 2 h prior to immunofluorescence. For Mps1 inhibition, mitotic cells were treated with 10 μM MPS1-IN-1 (Tocris). In the washout experiments, nocodazole or STLC was washed out by rinsing thrice with PBS followed by incubation with warm medium. Cells were fixed after 70 min. In RPE-1, to detect misaligned chromosomes, the cells were fixed 30 min after nocodazole washout. Microtubule stability assay was performed by triggering microtubule depolymerization with nocodazole treatment during 5, 15 and 30 min before fixation. For all live-cell experiments, drugs were added directly into the imaging medium and remained during the experiment.

### High-throughput Screening

HeLa cells were seeded in 96-well flat bottom plates (Capitol Scientific, Inc) at a density of 10,000 cells/100μL/well using Multidrop Combi Reagent Dispenser (Thermo-Scientific) and allowed to attach for 24 h. Cells were then blocked in mitosis with nocodazole for 3 h. After this period, 448 compounds from Cell Cycle/DNA Damage compound library (MedChemExpress) were added to cells at a final concentration of 10 μM using a Janus Automated Workstation (PerkinElmer) equipped with a pin tool (V&P Scientific). On each plate, DMSO and 10 μM etoposide were used as negative and positive controls, respectively. After 4 h, cells were fixed and stained for phospho-Histone H2AX (Ser139), phospho-Histone H3 (Ser10) and DAPI. The image acquisition was performed in an IN Cell Analyzer 2000 microscope (GE Healthcare) using a Nikon 20x/0.45 NA Plan Fluor objective, and nine fields of view (fov) per well were acquired. Image analysis was executed using the IN Cell Investigator software (GE Healthcare). Briefly, the image analysis workflow consists in the identification of the cell nuclei, from the DAPI channel. Then, the mean pixel intensity of the GFP and Texas-Red channels were measured for each cell. Spotfire software (TIBCO) was used to visualize and perform quality control of the data and to set the threshold to identify nuclei with positive GFP and Texas-Red signals. Cells with DNA damage were identified by the detected nuclei with positive GFP signal. Mitotic cells were defined by the cells with positive Texas-Red signal and mitotic cells with DNA damage were identified by overlap between positive GFP and Texas-Red signals. Hits were identified using a threshold of three standard deviations (3SD) above the mean of the negative control. The robustness of the screening assay was evaluated by the Z’-factor (Zhang et al., 1999):

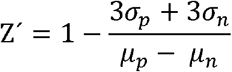

where μ_p_, μ_n_ and σ_p_ and σ_n_ represent mean and standard deviation of the positive and the negative controls, respectively. DMSO was used as a negative control and etoposide used as a positive control.

### Single-cell Backtracking

HeLa cells were imaged on a temperature controlled IN Cell Analyzer 2000 microscope (Nikon 20x/0.45 NA Plan Fluor objective). Cells were plated in 96-well flat bottom plates and incubated overnight in complete medium. Before live-cell imaging, cells were treated with nocodazole. The DNA-damaging compounds were added directly into the medium during the imaging. Images were acquired every 20 min. After 4 h treatment with DNA-damaging compounds, cells were fixed and stained for phospho-Histone H2A.X (Ser139), phospho-Histone H3 (Ser10) and DAPI. The image acquisition was performed in an IN Cell Analyzer 2000 microscope using Nikon 20x/0.45 NA Plan Fluor objective, and nine fov per well were acquired. A custom written Fiji/ImageJ (Schindelin et al., 2012) script was developed to automatically quantify the fluorescence intensity of γH2AX on mitotic cells. Importantly, after exposure to DNA-damaging compounds, only cells already in mitosis were analysed.

### Time-lapse Microscopy

HeLa cells were plated on a 12-well plate (Fisher Scientific) 24 h before imaging. Live-cell imaging experiments were performed at 37°C, in an atmosphere of 5% CO_2_, using a phase-contrast microscopy (Leica DMI6000; Leica Microsystems, 20x objective lens HCX PL FLUOTAR L. CORR. Ph1, 0.40 NA) equipped with a CCD camera (Hamamatsu FLASH4.0; Hamamatsu, Japan). To measure mitotic timing and to determine cell fate in mitotic arrested cells, nocodazole was added 1 h prior to acquisition. The other drugs were added directly into the medium during the imaging. The images were captured every 20 min for 72 h using LAS X software. In the washout experiment, HeLa cells stably expressing GFP-H2B/mRFP-α-tubulin were seeded 24 h prior to imaging on a μ-Plate 24-well Black (ibidi) and incubated overnight in phenol red free DMEM supplemented with 10% FBS and 25 mM HEPES (Gibco). Images were acquired every 10 min for 12 h using an IN Cell Analyzer 2000 microscope (GE Healthcare; Nikon 20x/0.45 NA Plan Fluor objective) at 37°C with 5% CO_2_. All images were analysed using Fiji/ImageJ.

### Immunofluorescence

Cells were fixed with 4% paraformaldehyde (Electron Microscopy Sciences) for 10 min and subsequently extracted in PBS-0.3% Triton (Sigma-Aldrich) for 10 min at room temperature (RT). After short washes in PBS-0.1% Triton, cells were incubated with blocking solution (10% FBS in PBS-0.1% Triton) for 1 h at RT and incubated with primary antibodies diluted in blocking solution overnight at 4°C. Then, cells were washed 3x with PBS-0.1% Triton and incubated with the respective secondary antibodies for 1 h at RT. DNA was counterstained with 1 μg/ml DAPI (Sigma-Aldrich). The coverslips were mounted using mounting medium (20 mM Tris pH 8, 0.5 mM N-propyl gallate, 90% glycerol) on glass slides. Mouse anti-phospho-Histone H2A.X Ser139 (1:2,000; Merck), rabbit anti-phospho-Histone H3 Ser10 (1:5,000; Cell Signaling Technology), mouse anti-Mad1 (1:500; Merck Millipore), guinea pig anti-CENP-C (1:1,000; MBL International Corporation), mouse anti–α-Tubulin clone B-512 (1:2,000:Sigma-Aldrich), rabbit anti–β-Tubulin (1:2,000; Abcam), rabbit anti-phospho-Aurora B Thr232 (1:1,000; Rockland Immunochemicals Inc.), rabbit anti-phospho-Aurora A Thr 288 (1:1,000; Cell Signaling Technology), mouse anti-Kid (1:500, a gift from S. Geley), rabbit anti-Kif4a (1,500; ThermoFisher Scientific) were used as primary antibodies, and Alexa Fluor 488, 568, and 647 (Invitrogen) were used as secondary antibodies (1:1,000). Images were acquired with AxioImager Z1 (63x, Plan oil differential interference contrast objective lens, 1.4 NA; all from Carl Zeiss) equipped with a charge-coupled device (CCD) camera (ORCA-R2; Hamamatsu Photonics) using the Zen software (Carl Zeiss). Forty-one z-planes with a 0.2 μm step covering the entire volume of the mitotic cell were collected. Autoquant X software (Media Cybernetics) was used for blind deconvolution. Images were analysed using Fiji/ImageJ and processed in Adobe Photoshop CS6 (Adobe Systems). Representative images were obtained through a maximum intensity projection of a deconvolved z stack and Adobe Illustrator CS6 (Adobe Systems) were used for panel assembly for publication.

### Fluorescence Quantification

For immunofluorescence quantitative measurements, all compared images were acquired using identical acquisition settings. Protein intensity was quantified using Fiji/ImageJ. Briefly, individual KTs were detected using CENP-C staining and a region of interest (ROI) delimitating the kinetochore on the focused Z plan was drawn. The mean fluorescence intensity of signal (pixel gray levels) of phospho-Aurora B at the inner centromere was measured on the focused z plan. The background was determined with 5 ROI drawn outside the KTs region and the average of these values were subtracted. Fluorescence intensity measurements were normalized to the CENP-C signals. Phospho-Aurora A levels at polar chromosomes was determined by drawing an elliptical ROI around the poles in sum-projected images. The background was determined with 5 ROI drawn outside the spindle poles region and the average of these values were subtracted. The microtubule depolymerization rate after nocodazole shock was determined by the proportion of total and soluble α-tubulin levels. The total α-tubulin intensity was measured by drawing a larger oval shaped ROI contained the entire cell in sum-projected images. The soluble α-tubulin levels were determined by drawing five smaller oval shaped ROI outside the chromosome region and the average of these values were calculated in sum-projected images. The fluorescence intensities were normalized to the level at time = 0 and represented as a function of time. The Kid and Kif4a levels were determined by drawing an elliptical ROI contained the entire cell in sum-projected images. The background was determined with 5 ROI outside the chromosome region and the average of these values were subtracted. All values were normalized to the average fluorescence levels of control cells.

### Western Blot

HeLa cells were resuspended and lysed in ice-cold NP-40 lysis buffer (20 mM HEPES-KOH pH 7.9, 1 mM EDTA, 1 mM EGTA, 150 mM NaCl, 0.5% NP-40, 10% Glycerol, 2 mM DTT) supplemented with a cocktail of protease and phosphatase inhibitors (Roche). The samples were kept on ice for 30 min and flash frozen in liquid nitrogen. To complement and increase lysis efficiency, cells were sonicated for 10 cycles of 30 sec “on” and 30 sec “off” at 4°C with Bioruptor (Diagenode) and the protein concentration was measured using Bradford protein assay (Thermo Fisher Scientific). Equal protein concentrations were denatured in Laemmli Buffer (62.5 mM Tris-HCl pH 6.8, 2% SDS, 10% glycerol, 5% 2-mercaptoethanol, 0.002% Bromophenol Blue) at 95 °C for 5 min, separated into an 8%-12% SDS-PAGE gels and transferred to a nitrocellulose Hybond-C membranes using a Trans-Blot Turbo Transfer System (Bio-Rad). Membranes were blocked with 5% milk in PBS-0.1% Tween-20 (Sigma-Aldrich; PBST) at RT during 1 h, and all primary antibodies were incubated overnight at 4°C with agitation. After five washes in PBST, the membranes were incubated with the secondary antibodies during 1 h at RT. After several washes with PBST, the detection was performed with Clarity Western ECL Blotting Substrate (Bio-Rad). Mouse anti-phospho-Histone H2A.X Ser139 (1:2,000; Merck) mouse anti-Mad1 (1:1,000; Merck), rabbit anti-Kif4a (1:1,000; ThermoFisher Scientific), mouse anti-Kid (1:1,000; a gift from S. Geley), mouse anti-MPM2 (1:1,000, Merck), mouse anti-GAPDH (1:50,000; Proteintech) were used as primary antibodies, and anti-mouse-HRP and anti-rabbit-HRP were used as secondary antibodies (1:5,000; Jackson ImmunoResearch Laboratories, Inc.).

### Statistical Analysis

Statistical analysis was performed using GraphPad Prism, version 8. All results presented in this manuscript were obtained from 2 independent experiments. All data represents mean ± S.D., and values were obtained across experiments or cells are indicated in the figure legend. D’Agostino-Pearson omnibus normality test was used to determine whether the data followed a normal distribution. p-Values were calculated with either t test (for data that followed a normal distribution) or Mann-Whitney Rank Sum test or a multiple comparison Kruskal-Wallis test (for data that did not follow a normal distribution). For the comparison of the single exponential fitting curve extra sum-of square F test was used. For each graph, n.s.= not significant, *p<0.05, **p≤0.01 ***p≤0.001 and ****p≤0.0001.

## Supplemental Figures Legends

**Figure S1 (related to Figure 1). Interphase HeLa cells acquire DNA damage. (A)** Percentage of interphase HeLa cells with γH2AX positive after 4 h treatment with chemical library of 448 cell cycle inhibitors/DNA-damaging compounds. DMEM and DMSO (green dots) and etoposide (blue dots) were used as negative and positive controls. Each dot represents a compound. The thresholds (dash lines) were defined as mean (Avg) plus or minus three times standard deviation (3SD) from negative controls. Compounds above the threshold were consider hits (yellow dots). **(B)** Western blot analysis of γH2AX levels from mitotic HeLa cell extracts. Mitotic extracts were derived from isolated mitotic cells treated with DMSO or the selected mitotic DNA-damaging compounds. GAPDH was used as loading control. Approximate molecular weights are shown on the left. The various classes of DNA-damaging compounds are represented by different colours: DNA-alkylating agents (in green), DNA-intercalating agents (in yellow), DNA-crosslinking agents (in purple), DNA adduct-inducing agents (in orange) and agents that generate single- or double-strand breaks (in blue).

**Figure S2 (related to Figure 2). Long-term mitotic DNA damage affects mitotic fidelity.** Detailed description of cell fates listed as “other” shown in figure 2C.

**Figure S3 (related to Figure 2). Long-term mitotic DNA damage persists throughout mitosis. (A)** Representative immunofluorescence images of HeLa cells treated with nocodazole and DMSO during 4 h, followed by nocodazole washout and fixation after 70 minutes. Cells were immunostained with anti-CENP-C (gray), anti-β-tubulin antibody (magneta), anti-γH2AX antibody (green) and DAPI (blue). Scale bar = 5 μm. **(B)** Representative immunofluorescence images of HeLa cells treated with nocodazole and etoposide during 4 h, followed by nocodazole washout and fixation after 70 minutes. Cells were immunostained with anti-CENP-C (gray), anti-β-tubulin antibody (magneta), anti-γH2AX antibody (green) and DAPI (blue). Scale bar = 5 μm.

**Figure S4 (related to Figure 4). Long-term mitotic DNA damage also leads to the formation of stable monotelic attachments on polar chromosomes after mitotic arrest/washout with the kinesin-5 inhibitor. (A)** Representative immunofluorescence images of HeLa cells treated with S-trityl-L-cysteine (STLC) and either DMSO or etoposide during 4 h, followed by STLC washout and fixation after 70 minutes. Cells were immunostained with anti-CENP-C (gray), anti-β-tubulin antibody (magneta), anti-Mad1 antibody (green) and DAPI (blue). Scale bar = 5 μm. **(B)** Quantification of Mad1 detection on kinetochores pairs from polar chromosomes under different conditions presented in A.

**Figure S5 (related to Figure 5). Etoposide induces mitotic DNA damage in U2OS cells, Related to Figure 5.** Representative immunofluorescence images of prometaphase arrest U2OS cells treated with DMSO or etoposide during 4 h after nocodazole treatment, with the indicated antibodies. Scale bar = 5 μm.

**Figure S6 (related to Figure 5). Long-term mitotic DNA damage increases kinetochore-microtubule attachment stability in U2OS cells. (A)** Representative immunofluorescence images of U2OS cells treated with nocodazole and either DMSO or etoposide during 4 h, followed by nocodazole washout and fixation after 70 minutes. Cells were immunostained with anti-CENP-C (gray), anti-β-tubulin antibody (magneta), anti-Mad1 antibody (green) and DAPI (blue). Scale bar = 5 μm. **(B)** Quantification of Mad1 detection on kinetochores pairs from polar chromosomes under different conditions presented in A. **(C)** Representative immunofluorescence images of U2OS cells treated with nocodazole and either DMSO or etoposide during 4 h, followed by nocodazole washout. After 70 minutes, a nocodazole shock was applied and the cells were fixed after 5, 15 and 30 minutes. Cells were immunostained with anti-β-tubulin antibody (inverted grey scale) and DAPI (blue). Scale bar = 5 μm. Taxol was used as a positive control. **(D)** Normalized ratio of total α-tubulin/soluble α-tubulin fluorescence intensity over time. The kinetics of fluorescence decay after nocodazole shock were determined as a readout of resistance to microtubule depolymerization. Whole lines show single exponential fitting curve. Each data point represents the mean ± SD; * p<0.05 relative to control, extra sum-of-square F test.

**Figure S7 (related to Figure 5). Long-term mitotic DNA damage does not affect kinetochore-microtubule attachment stability in RPE-1 cells. (A)** Representative immunofluorescence images of prometaphase arrest RPE-1 cells treated with DMSO or etoposide during 4 h after nocodazole treatment, with the indicated antibodies. Scale bar = 5 μm. **(B)** Representative immunofluorescence images of RPE-1 cells treated with nocodazole and either DMSO or etoposide during 4 h, followed by nocodazole washout. After 30 minutes, a nocodazole shock was applied and the cells were fixed after 5, 15 and 30 minutes. Cells were immunostained with anti-β-tubulin antibody (inverted grey scale) and DAPI (blue). Scale bar = 5 μm. Taxol was used as a positive control. **(C)** Normalized ratio of total α-tubulin/soluble α-tubulin fluorescence intensity over time. The kinetics of fluorescence decay after nocodazole shock were determined as a readout of resistance to microtubule depolymerization. Whole lines show single exponential fitting curve. Each data point represents the mean ± SD; n.s.: not significant relative to control, extra sum-of-square F test.

**Figure S8 (related to Figure 6). Long-term mitotic DNA damage does not affect Aurora A and Aurora B activity on polar chromosomes. (A)** Representative immunofluorescence images of HeLa cells treated with nocodazole and either DMSO or etoposide during 4 h, followed by nocodazole washout and fixation after 70 minutes. Cells were immunostained with anti-CENP-C (gray), anti-α-tubulin antibody (magneta), anti-phospho-Aurora A Thr 288 antibody (green) and DAPI (blue). Scale bar = 5 μm. **(B)** Normalized fluorescence intensity of pAurora A at the poles. Each dot represents an individual cell. (DMSO, 1.00 ± 0.28, n = 20; Etoposide, 0.98 ± 0.27, n = 20; mean ± SD; n.s.: not significant relative to control, t test). **(C)** Representative immunofluorescence images of HeLa cells treated with nocodazole and either DMSO or etoposide during 4 h, followed by nocodazole washout and fixation after 70 minutes. Cells were immunostained with anti-CENP-C (grey), anti-α-tubulin antibody (magneta), anti-phospho-Aurora B Thr 232 antibody (green) and DAPI (blue). Scale bar = 5 μm. **(D)** Relative fluorescence intensity of pAurora B/CENP-C at inner centromere in the unaligned to aligned chromosome. Each dot represents an individual kinetochore. (DMSO, 1.46 ± 0.41, n = 20; Etoposide, 1.46 ± 0.32, n = 25; mean ± SD; n.s.: not significant relative to control, Mann-Whitney Rank Sum test).

## Acknowledgments

We would like to thank all colleagues that kindly provided reagents used in this study and the support of the i3S Scientific Platforms: BioSciences Screening and Advanced Light Microscopy, members of the national infrastructure PT-OPENSCREEN (NORTE-01-0145-FEDER-085468) and PPBI – Portuguese Platform of Bioimaging (PPBI-POCI-01-0145-FEDER-022122). M. Novais-Cruz holds a PhD fellowship (SFRH/BD/117063/2016 and COVID/BD/151730/2021) and C. Ferrás was supported by an Investigator starting grant (IF/00765/2014) from Fundação para a Ciência e a Tecnologia of Portugal. This work was funded by FCT (IF/00765/2014/CP1241/CT0003 to C. Ferrás and PTDC/MED-ONC/3479/2020 to H. Maiato), by NORTE 2020 under the PORTUGAL 2020 Partnership Agreement through the European Regional Development Fund (NORTE-01-0145-FEDER-000029 to C. Ferrás and NORTE-01-0145-FEDER-000051 to H. Maiato), as well as by the European Research Council (ERC) consolidator grant CODECHECK, under the European Union’s Horizon 2020 research and innovation programme (grant agreement 681443) and a La Caixa Health Research Grant (LCF/PR/HR21/52410025) to H. Maiato.

## Author contributions

Methodology (MNC, AP, MS, AFM, CF); Investigation, Formal Analysis and Validation (MNC, AP, MS, AFM, HM, CF); Visualization (MNC, AP, MS, AFM, HM, CF); Writing – Original Draft (MNC, HM); Writing – Review and Editing (MNC, AP, MS, AFM, HM, CF); Conceptualization, Supervision, Project Administration and Funding acquisition (HM, CF).

