## Supplementary figures and images for "Long-term mitotic DNA damage promotes chromokinesin-mediated missegregation of polar chromosomes in cancer cells"

### Figure S1

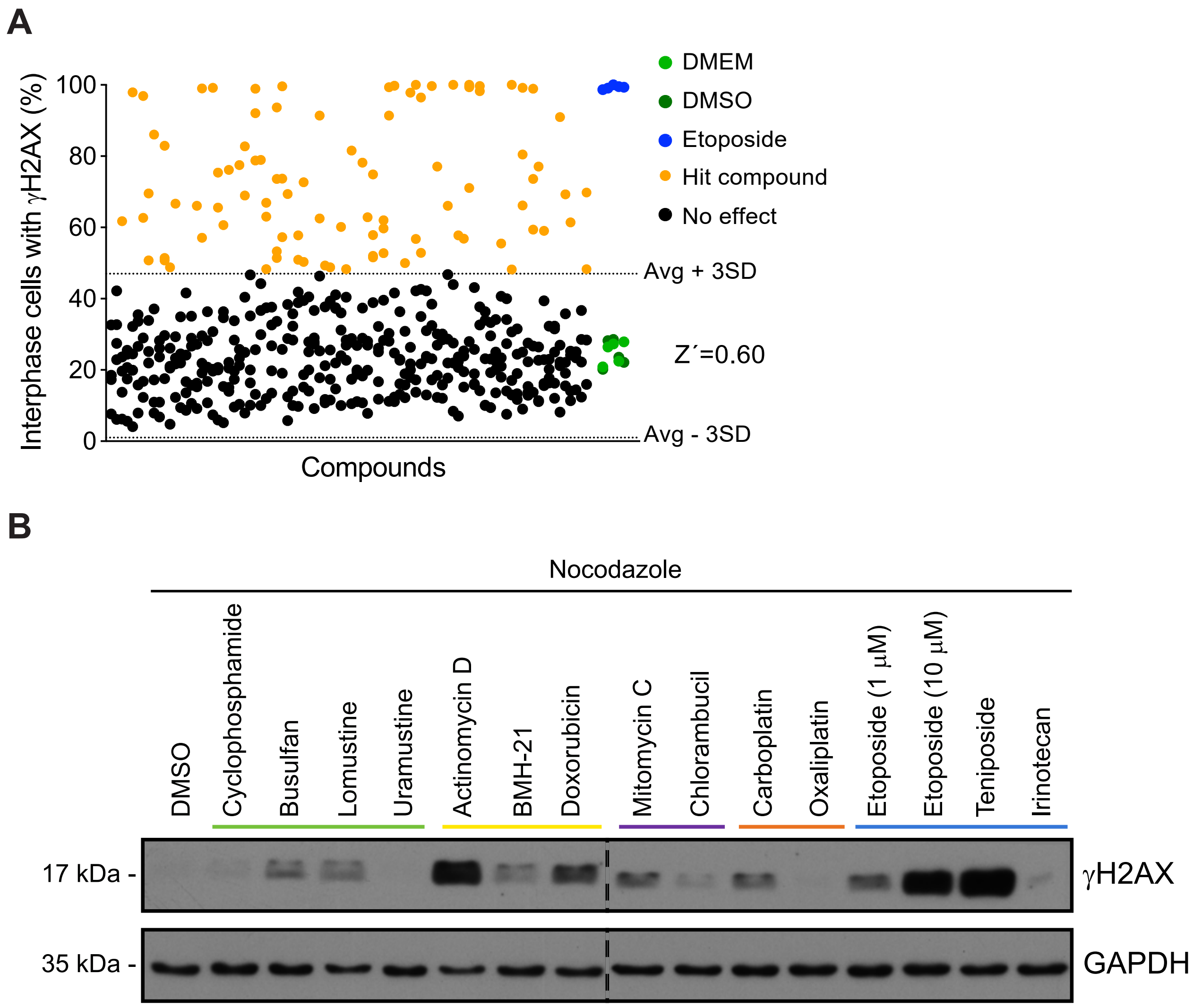

### Figure S2

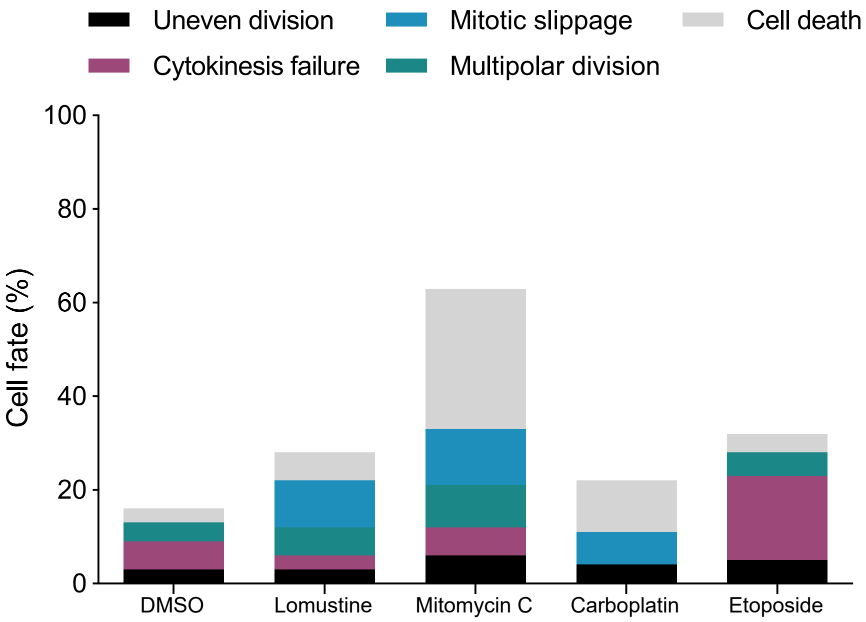

### Figure S3

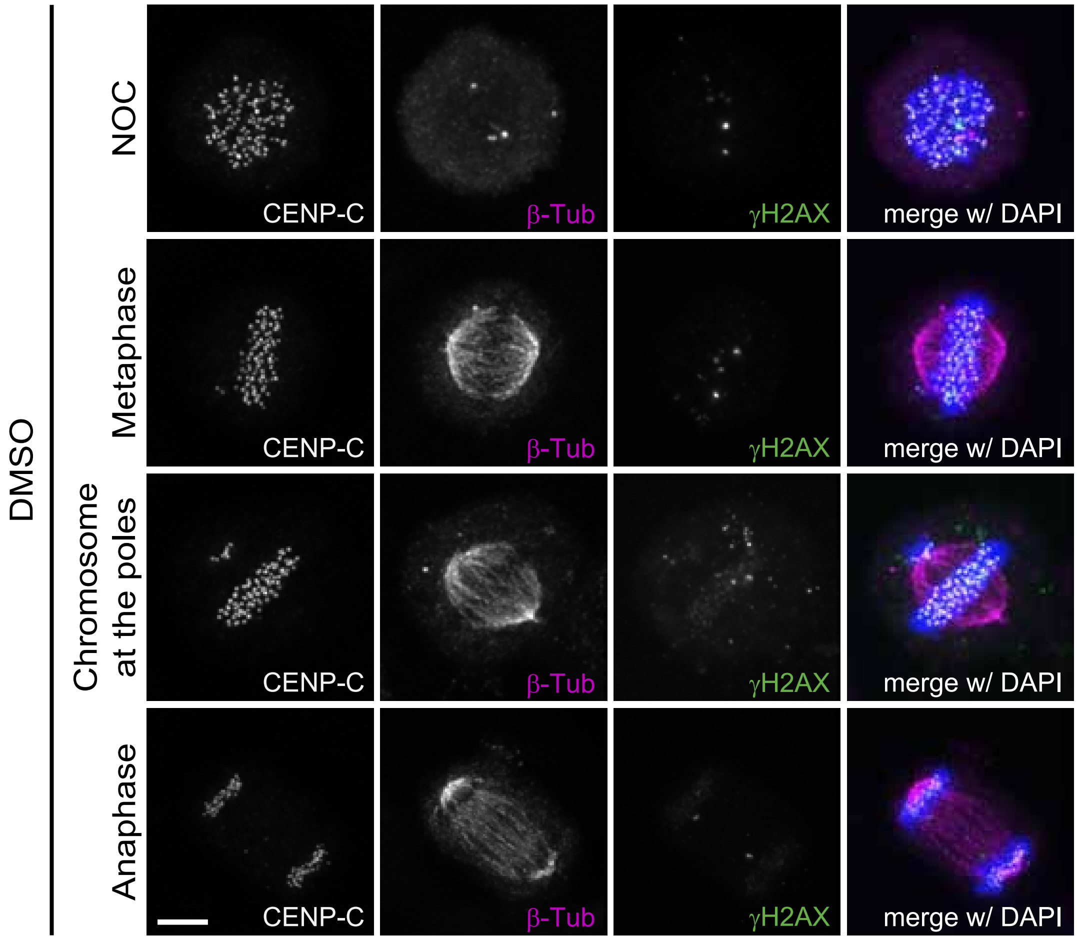

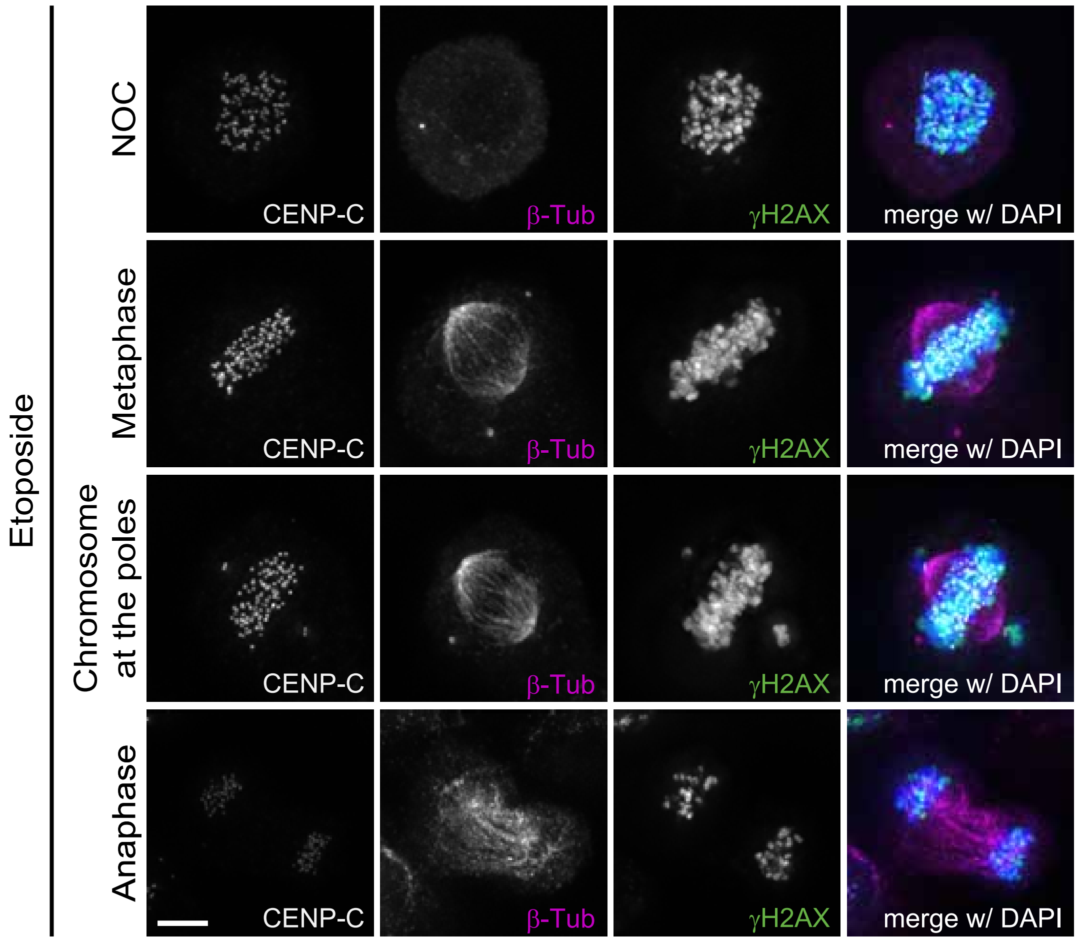

### Figure S4

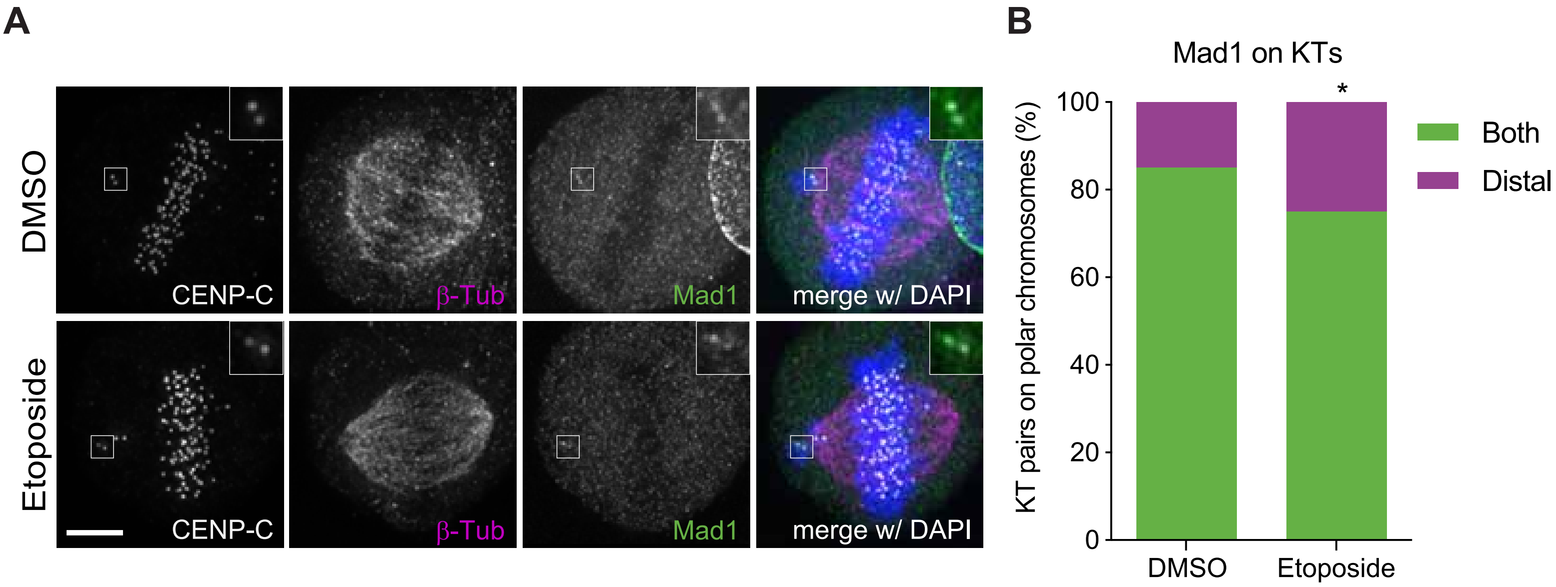

### Figure S5

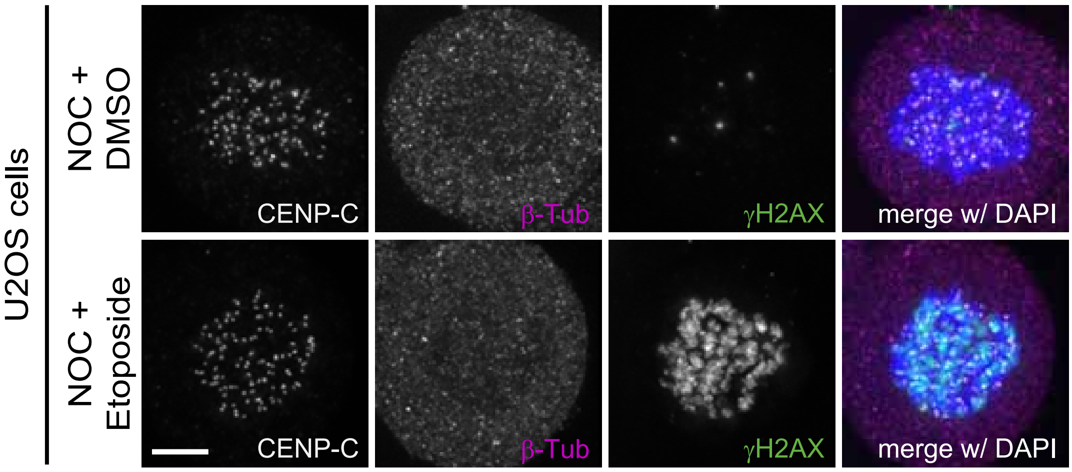

### Figure S6

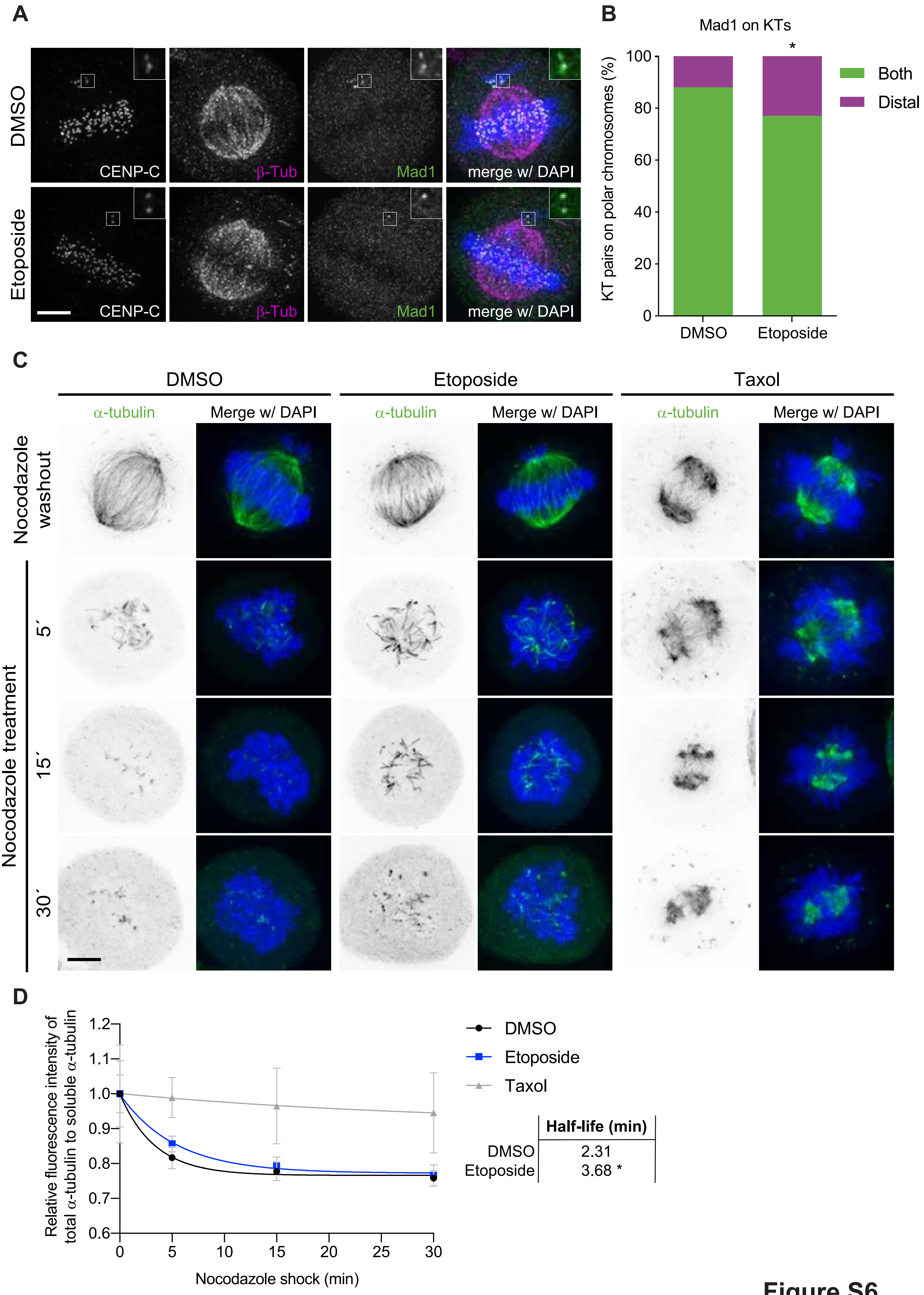


| | Half-life (min) |
|---|---|
| DMSO | 2.31 |
| Etoposide | 3.68 * |

**Figure S6**

### Figure S7

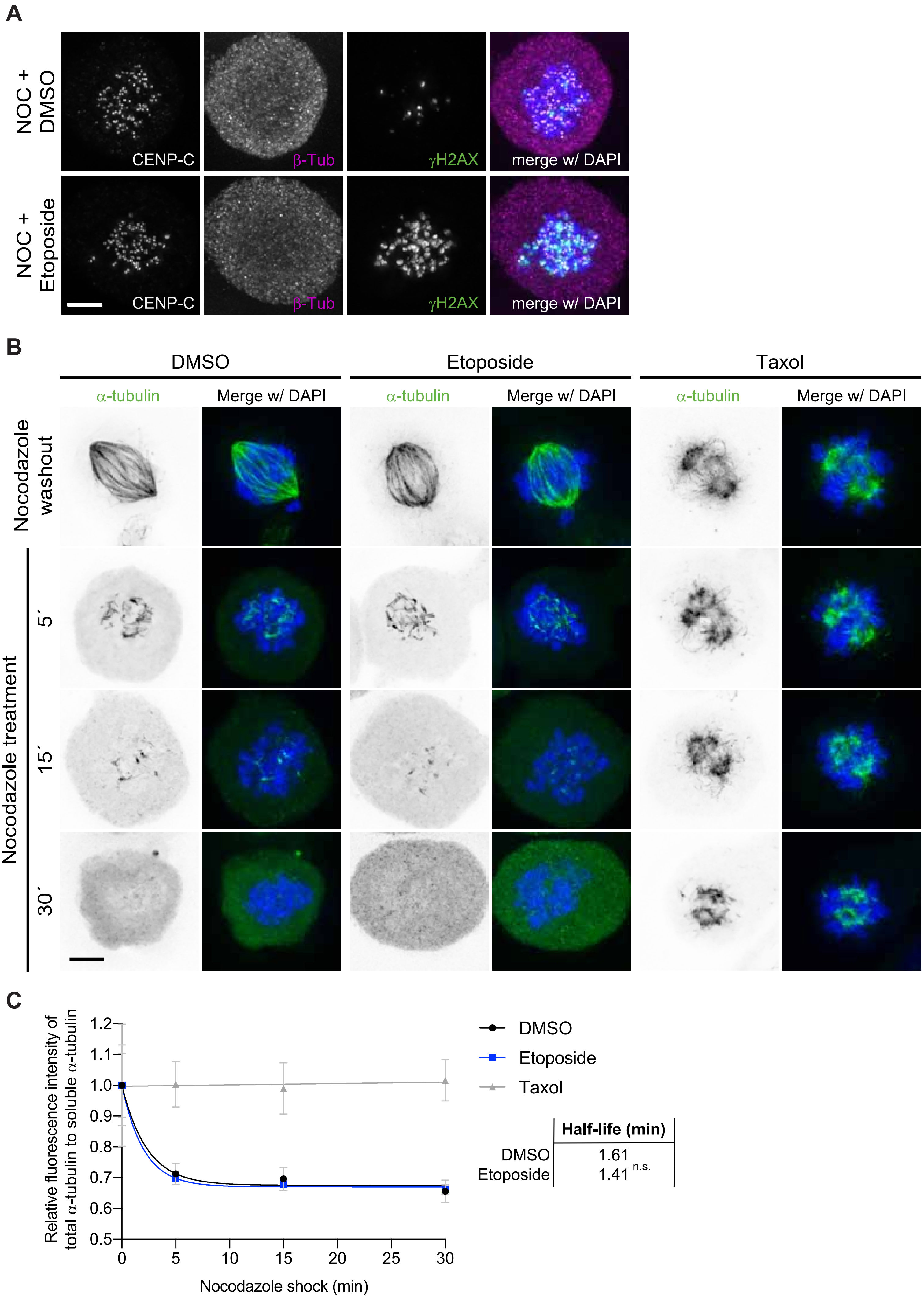

### Figure S8

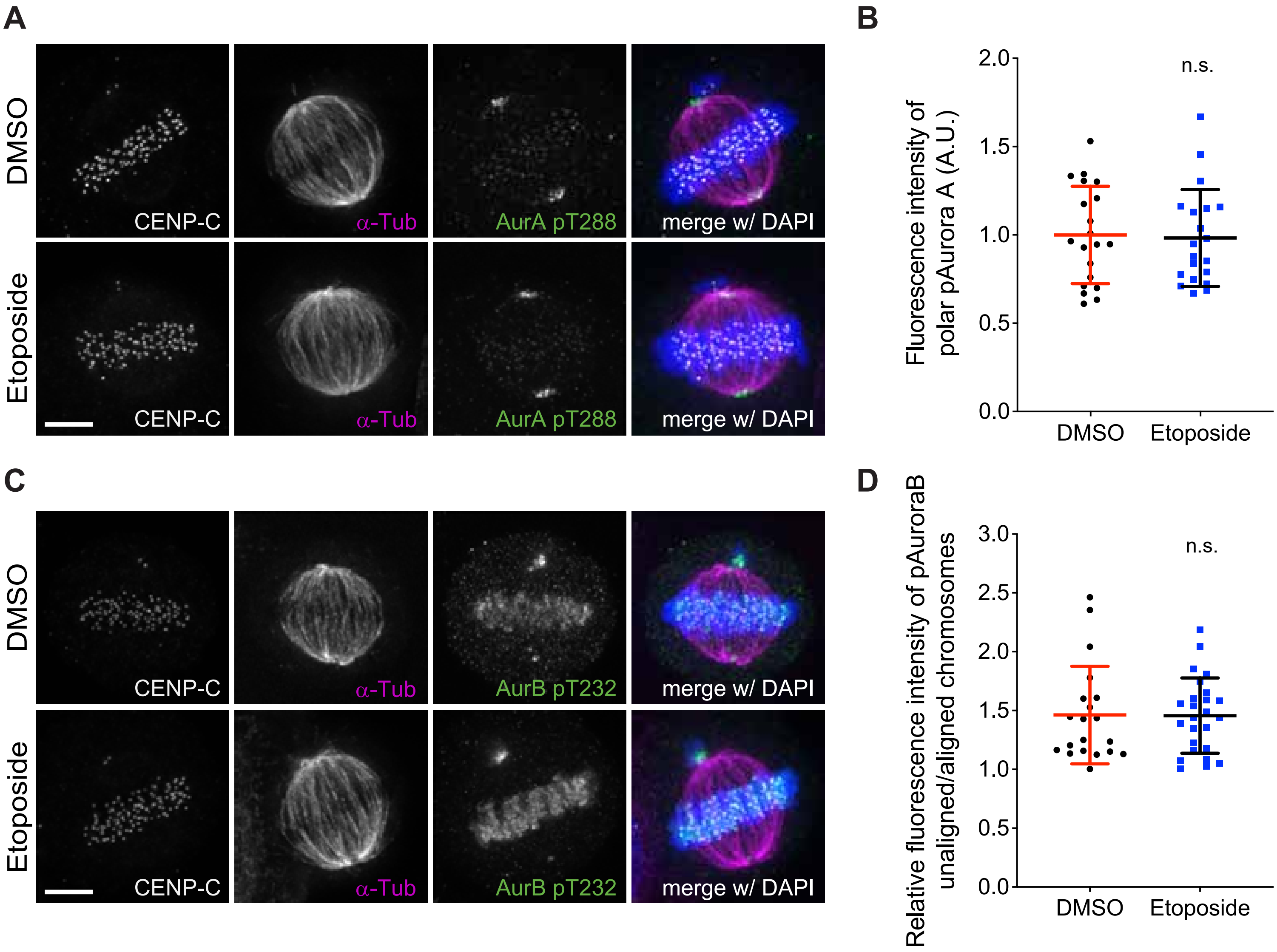
